# CatESO: Differentiable Enzyme Sequence Optimization Guided by Substrate-Aware kcat Prediction

**DOI:** 10.64898/2026.07.04.736506

**Authors:** Zhenjia Gan, Yuzhi Xu, Junde Xu, Zhihao Wu, Juping Huang, Jiabin Yin, Guangyong Chen, John Z.H. Zhang

## Abstract

Enzymes drive biological chemistry and offer greener routes to chemicals, materials and medicines, yet their broader use as biocatalysts is often limited by insufficient catalytic turnover. Improving turnover is hard: measured rate constants are scarce and protein sequence space is vast. Deep-learning models now predict the turnover number, *k*_cat_, with growing accuracy, but they are typically applied after sequence generation to score or filter candidates, which separates the kinetic objective from the design itself. To bridge the gap between sequence generation and kinetic evaluation, we introduce CatESO, a differentiable sequence optimizer that enables direct, gradient-guided design of substrate-specific catalytic turnover. By backpropagating through a cross-modal *k*_cat_ predictor under continuous sequence relaxation, CatESO co-optimizes predicted catalytic activity, evolutionary plausibility and structural integrity in one end-to-end framework, using ESM-2 and ESMFold to keep designs evolutionarily plausible and foldable. Across seven stringent out-of-distribution enzymes spanning EC classes 1–7, CatESO raised model-predicted *k*_cat_ for the vast majority of designs, with a median predicted fold change of 1.52 while every variant retained a pLDDT above 70. Against RFdiffusion3–LigandMPNN pipeline and ZymCtrl, CatESO struck a better balance between predicted activity and structural confidence. By making substrate-conditioned kinetic objectives differentiable, CatESO carries differentiable protein design beyond structure- and binding-centred goals to enzyme catalytic function, giving a general route to function-oriented enzyme engineering.

## Introduction

Through evolution, nature has generated a vast repertoire of enzymes, protein catalysts that accelerate the majority of chemical transformations required for biological systems to survive across diverse environments and are abundant in all living cells.^1–5^ To repurpose these catalysts, enzyme engineering has long drawn on three complementary strategies:^6–10^ rational design introduces targeted point mutations at residues identified from experimental structures or homology models; ^11–14^ directed evolution improves performance through iterative rounds of random mutagenesis and screening;^15–20^ and semi-rational design combines structural, sequence and mechanistic information to build functionally enriched variant libraries.^21–25^ A common target of these efforts is the turnover number, *k*_cat_, which measures how fast an enzyme converts substrate to product under saturating conditions.^26–30^ Yet measured *k*_cat_ values are scarce, and sequence space is so large that experiments can sample only a minute fraction of possible variants^31–34^ (Fig. 1A). Each of the three strategies therefore tends to search a limited, mostly local region of the sequence–fitness landscape, and distant high-performing variants stay hard to reach.^35–40^

**Figure 1:**
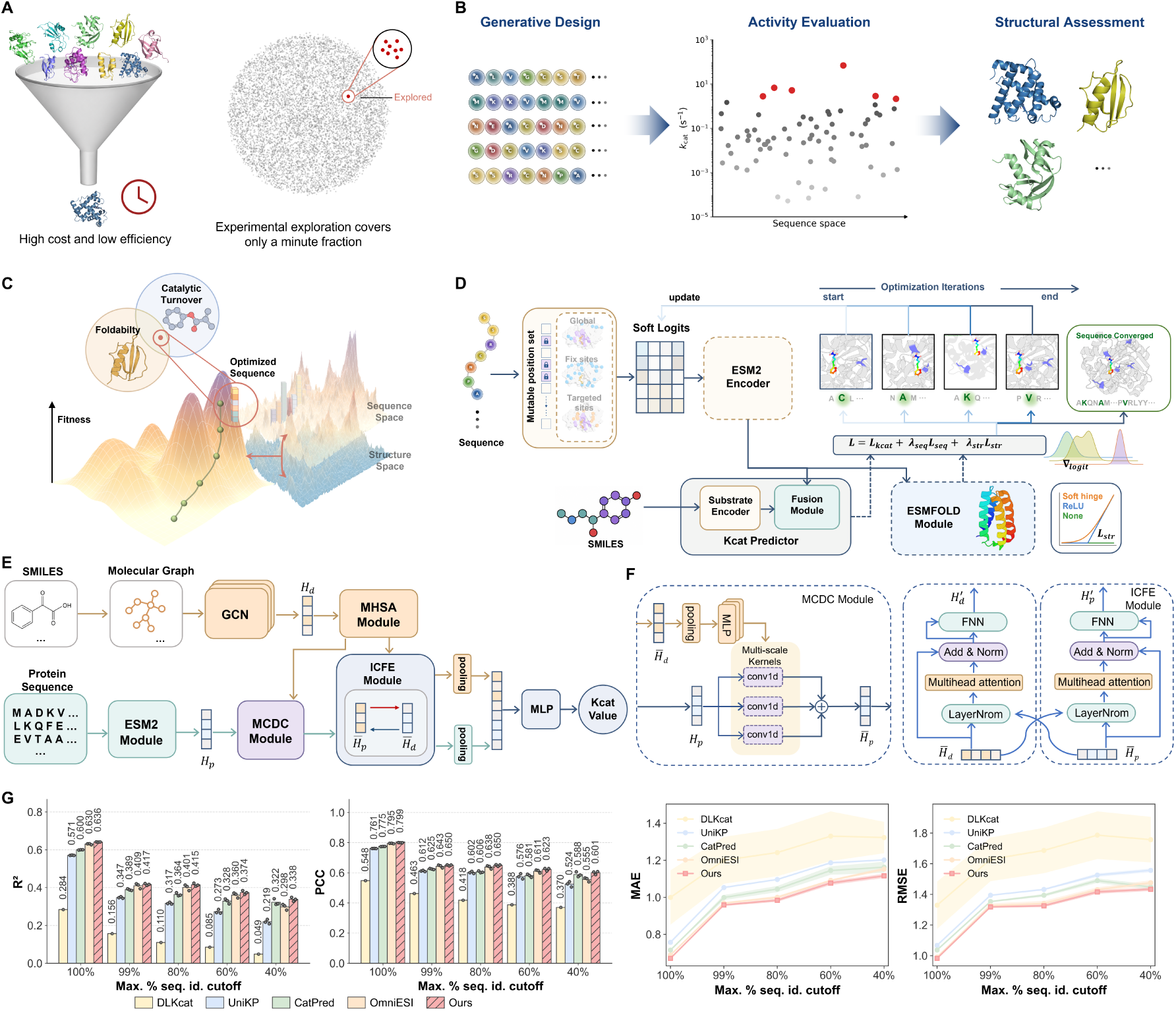
CatESO: a differentiable framework for substrate-aware *k*_cat_ prediction and enzyme sequence optimization. (A) Conventional enzyme engineering is experimentally intensive and explores only a sparse, largely local fraction of protein sequence space. (B) Current *k*_cat_ predictors are used to score and rank candidates once they exist, leaving prediction decoupled from the design step. (C) Schematic of gradient-guided optimization, in which gradients from a differentiable activity predictor flow back to a continuous sequence representation and drive it toward higher predicted activity. (D) Overview of the CatESO workflow. Sequence logits are optimized to increase the predicted *k*_cat_, while ESM-2-derived sequence plausibility and ESMFold-derived structural confidence regularize the search; the relaxed sequence representation is gradually discretized during optimization. (E) Architecture of the CatESO *k*_cat_ predictor. Substrate and enzyme inputs are encoded separately, coupled by the MHSA, (F) MCDC and ICFE modules, and mapped by an MLP to log_10_(*k*_cat_). (G) Benchmark comparison on CatPred-DB against DLKcat, UniKP, CatPred and OmniESI across five maximum sequence-identity cutoffs. Bar plots show PCC and *R*^2^; line charts show MAE and RMSE.

AI-driven methods have begun to address this limitation along two largely separate ways. The first treats enzyme kinetic prediction as a multimodal learning problem, fusing protein sequence or structure with substrate-derived molecular features. DLKcat,^35^ UniKP,^41^ Cat-Pred^42^ and OmniESI^43^ successively added deep sequence encoders, protein language models, molecular graphs and enzyme–substrate feature fusion, and each gain in *k*_cat_ accuracy points to the same conclusion: catalytic activity is read out from the enzyme–substrate pair, not from either partner alone^35,44,45^ (Fig. 1B). The second pursues generative protein design, which now goes beyond fixed backbones to design sequences, backbones, interfaces and catalytic sites under user-defined constraints.^46–54^ Here RFdiffusion2,^55^ LigandMPNN,^56^ ZymCTRL,^57^ EnzymeFlow^58^ and GENzyme^59^ have moved enzyme design from EC-guided generation toward reaction-aware modelling of catalytic architectures.^60^ These two lines rarely meet. Most generate-then-screen workflows use predicted activity only as a terminal filter: candidates are designed first and scored afterwards, so turnover does not guide the search.^61–63^ An accurate *k*_cat_ predictor is therefore the natural lever for closing this gap.^35^ In the idealized limit, a perfect predictor would reduce the activity side of design to an oracle-ranking problem—score candidate sequences and retain those with the highest predicted turnover, in the spirit of the “reward is enough” view from reinforcement learning.^64^ But sequence space is far too large to score exhaustively.^65^ The value of a strong predictor is not that it enables brute-force screening,^65,66^ but that it localizes the search: it points to a small region likely to contain high-turnover variants,^67^ and its gradient shows how to move a sequence toward higher predicted turnover.^68,69^ The search can then optimize within that region directly instead of sweeping the whole space.

This gradient-guided view has a concrete precedent: pretrained biomolecular models^70^ can be inverted for sequence generation. Early deep-network hallucination repurposed structure-prediction networks such as trRosetta for de novo protein design, optimizing random sequences against model-derived structural objectives.^71,72^ Relaxed-sequence and AlphaFold-based implementations later made this inversion differentiable, letting gradients from predictive models guide sequence search across larger proteins and more demanding design settings.^73^ Differentiability alone, however, does not guarantee realistic designs. Optimizing directly against a predictor’s own confidence is the same procedure that produces adversarial examples:^74,75^ gradient ascent can exploit model imperfections and return high-confidence sequences that are biophysically unrealistic or poorly expressible. Working pipelines therefore pair model inversion with safeguards, among them sequence redesign with Protein-MPNN or its soluble variant, ensembles of prediction models, progressive discretization, independent structure re-prediction and Rosetta or interface filtering before experimental validation.^72,73,76^ BindCraft makes this trade-off concrete. It backpropagates through AlphaFold2/AF2-multimer to co-optimize binder sequences, structures and interfaces, then redesigns surfaces with soluble ProteinMPNN and applies stringent AF2 and Rosetta filtering, producing experimentally validated binders—many with nanomolar affinity—across diverse targets.^76^ Predictive models can thus serve as generative design engines rather than after-the-fact scoring functions, but only when the differentiable objective sits inside a constraint-aware loop. For enzyme engineering, the question follows directly: can a substrate-conditioned kinetic objective be made differentiable and used to steer sequence search itself, while structural sequence-manifold and active-site constraints keep that search from drifting toward unrealistic or catalytically incompatible variants?

We close this loop with CatESO, a catalytic-turnover-guided enzyme sequence optimizer that makes a substrate-conditioned kinetic objective differentiable and uses it to steer sequence search directly, under constraints that keep designs evolutionarily and structurally realistic.^77,78^ Such an optimizer is only as trustworthy as the gradient behind it, so we first train an accurate substrate-aware *k*_cat_ predictor that couples ESM-2-derived protein representations with a graph-based encoding of the substrate and reads turnover off the enzyme–substrate pair. On the curated CatPred-DB benchmark it surpasses DLKcat, ^35^ UniKP,^41^ CatPred^42^ and OmniESI,^43^ most clearly on low-identity sequences—the regime an optimizer must enter—so its gradient stays reliable where the search travels. CatESO then inverts this predictor into a generator. For a given enzyme–substrate pair, it represents mutable residues as differentiable soft sequences through a Gumbel–Softmax relaxation,^78–81^ optimizing mutation probabilities continuously instead of searching over discrete variants (Fig.1C). ESM-2^82^ then encodes these relaxed sequences and keeps the search within an evolutionarily plausible region of sequence space. Gradients from an ensemble of the trained kinetic predictors, together with differentiable constraints from ESM-2 and a structure predictor, flow back to the shared soft-sequence variables, so predicted activity, sequence plausibility and folding confidence jointly shape residue preferences during generation (Fig.1D). Conventional generate–predict–filter pipelines keep these criteria in separate stages; CatESO folds them into a single optimization trajectory that raises predicted turnover while holding the search on the manifold ESM-2 supports and away from variants unlikely to fold. CatESO thereby carries differentiable design past its usual structure- and binding-centred goals to enzyme catalytic function.

## Results and discussion

### Substrate-conditioned kinetic modelling improves *k*_cat_ prediction

The differentiable design loop in Figure 1D rests on the *k*_cat_ predictor that supplies the optimization gradient. The same predictor plays two roles: used only after sequence generation it filters generated candidates, whereas embedded within the design process it becomes the objective that shapes the search. Its errors propagate through the optimization trajectory, and how informative its gradients are determines which regions of sequence space are explored. The predictor must therefore satisfy two requirements at once: it must estimate catalytic turnover accurately, and it must give a smooth enough objective to guide sequence optimization rather than only rank candidates after they are generated.

CatESO was designed to meet these requirements. It estimates log_10_(*k*_cat_) from an enzyme sequence and its substrate or reactant set (Fig. 1E), combining ESM-2-derived protein representations with graph-based molecular encodings so that turnover is inferred from the enzyme–substrate pair rather than from either component alone. The MCDC and ICFE modules condition residue-level protein features on the reactant context and let enzyme and molecular representations exchange information in both directions (Fig. 1F). The resulting interaction fingerprint passes to a regression head, and an ensemble of independently trained predictors provides the estimate used during optimization. CatESO is therefore both a substrate-aware kinetic predictor and a differentiable objective for guiding enzyme sequence design.

We evaluated CatESO on the curated CatPred-DB benchmark using the original data partitions, which allows direct comparison with DLKcat, UniKP, CatPred and OmniESI (Fig. 1G, Table. S1). We measured performance at five maximum sequence-identity cutoffs: 100%, 99%, 80%, 60% and 40%. Lowering the cutoff reduces the similarity between training and test enzymes, so each split tests generalization beyond close homologues more strictly than the last. CatESO was the strongest method at every cutoff, with *R*^2^ values of 0.636, 0.417, 0.415, 0.374 and 0.338 and the lowest MAE values of 0.670, 0.959, 0.984, 1.077 and 1.116 across the same splits. PCC and RMSE followed the same pattern, so the gain holds for both correlation- and error-based metrics rather than for one alone.

Aggregate performance can obscure heterogeneous behaviour across enzyme families or sequence lengths. We therefore stratified the benchmark by EC class and protein length (Fig. 2A, Table. S2, S3). Across the three length ranges (*l* ≤ 300, 300 *< l* ≤ 700 and *l >* 700) and most of the seven EC classes, CatESO achieved the highest subset-level *R*^2^ and PCC, while remaining competitive in the few categories where another method performed best. More importantly, CatESO showed substantially less performance fluctuation across subsets. In contrast, CatPred, UniKP and DLKcat varied markedly between enzyme groups, with strong performance in some categories but clear degradation in others. Thus, CatESO’s overall advantage is not driven by a few favourable or well-represented subsets, but reflects more uniform accuracy across diverse reaction chemistries and protein sizes.

**Figure 2:**
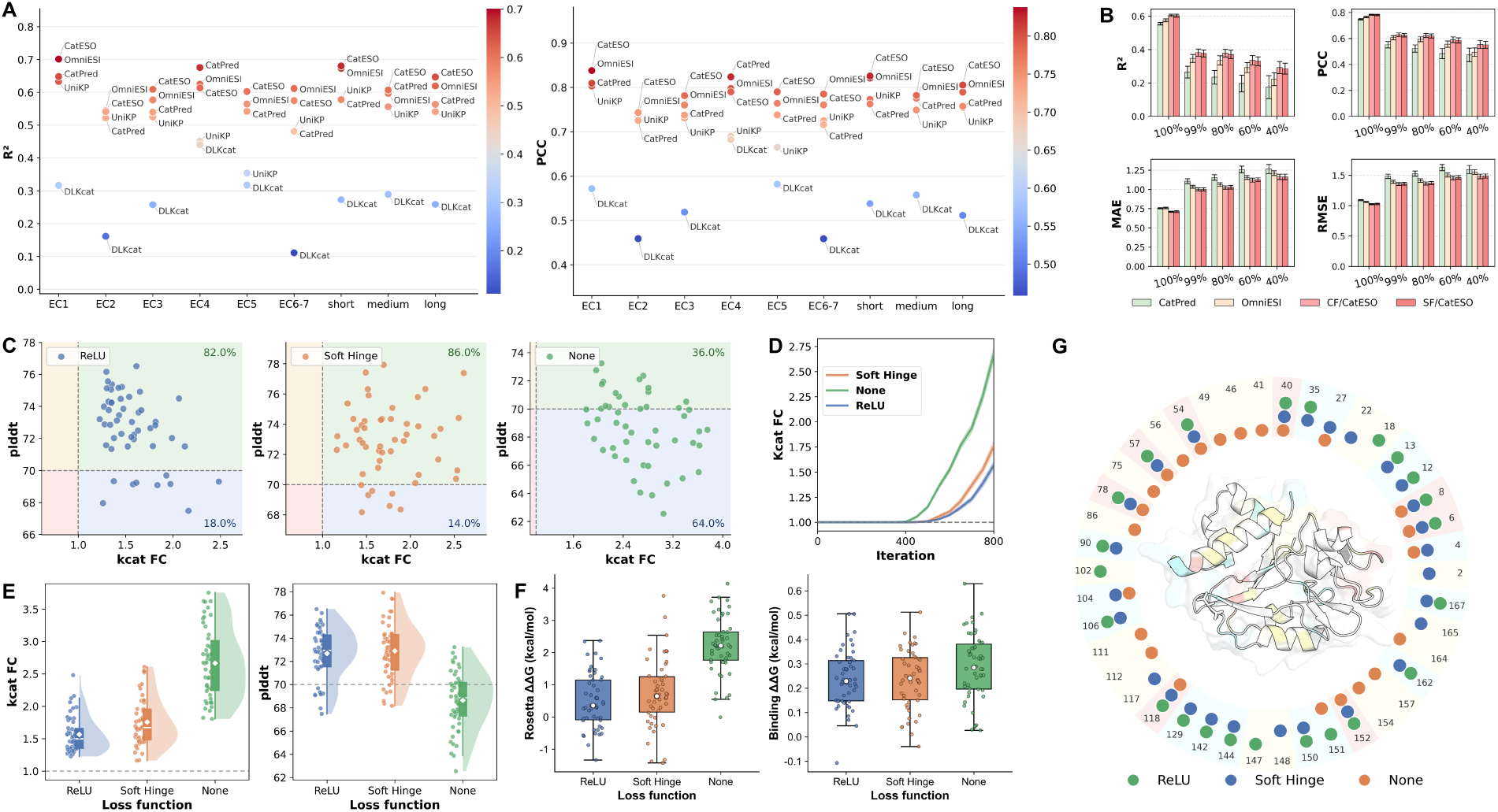
Predictor benchmarking and structure-constrained sequence optimization. (A) Stratified performance across EC classes and protein-length bins. Each point is the subset-level *R*^2^ or PCC of one model, evaluated over seven EC classes and three length ranges. (B) Single-model comparison of the predictors that make up the CatPred, OmniESI and CatESO ensembles. CatESO is shown with two prediction heads, concat fusion (CF) and simple fusion (SF). *R*^2^, PCC, MAE and RMSE are reported across the five sequence-identity cutoffs. (C) Fifty optimized Q9P9E3 sequences under a ReLU, soft-hinge, or no structural penalty. *k*_cat_ fold change is predicted by CatESO and pLDDT by ESMFold; dashed lines mark *k*_cat_ FC = 1 and pLDDT = 70. (D) Predicted *k*_cat_ fold-change trajectories during optimization, showing the trade-off between maximizing activity and preserving structural confidence. (E) Final *k*_cat_ fold-change and pLDDT distributions for the three penalty settings. (F) Final Rosetta ΔΔ*G* and binding ΔΔ*G* distributions for the three penalty settings. (G) Positions of CatESO-selected mutations under the three penalty settings, mapped around the Q9P9E3 structure.

Finally, we asked whether the improvement was just model averaging. We compared the individual predictors behind the CatPred, OmniESI and CatESO ensembles and tested CatESO with both concat fusion (CF) and simple fusion (SF) prediction heads (Fig. 2B, Table. S4). A single CatESO predictor still beat the baselines at all five cutoffs and on *R*^2^, PCC, MAE and RMSE (Table. S1). The gain thus comes mainly from the substrate-aware interaction architecture, not from ensembling. The advantage also held under the strictest low-identity splits, where test enzymes are most distant from the training set. This is the regime a differentiable enzyme designer must enter once it moves beyond local mutational neighbourhoods, which suggests that CatESO offers not only a stronger predictor but a more reliable gradient source for searching remote sequence space. This reliability is consistent with its architecture: relative to OmniESI, CatESO is overall shallower, chiefly in its feature-fusion modules, which shortens the gradient path back to the sequence logits. Its wider projection and interaction modules add parameters, but the network still backpropagates the kinetic objective more directly and stably.

### Structural constraints prevent adversarial drift in differentiable enzyme design

The trained predictor gives reliable gradients, so we next turned it toward enzyme sequence design directly (Fig. 1D). CatESO turns the substrate-conditioned *k*_cat_ predictor into a differentiable design objective: for a given reference enzyme and target substrate set, mutable residues are represented by continuous logits, and gradients from the ensemble-predicted log_10_(*k*_cat_) objective are propagated back to these logits to bias the sequence search toward higher predicted turnover. This allows optimization directly in sequence space, rather than retrospective scoring of pre-generated variants. It also exposes a familiar risk of gradient-based model exploitation: optimizing an input against a predictor’s own output can produce adversarial solutions that score highly while violating the physical constraints of the underlying system.^74,75^ CatESO therefore couples the activity objective with sequence- and structure-aware constraints.^76^We began with a representative enzyme to probe whether these constraints are needed to keep differentiable optimization within a realistic region of sequence space.

We selected Q9P9E3, identified as described in the Methods, and optimized its sequence under three structural-regularization settings while holding all other hyperparameters fixed: a ReLU penalty, a soft-hinge penalty, or no structural penalty (Fig. 2C). For each setting we generated 50 sequences and scored them by CatESO-predicted *k*_cat_ fold change (*k*_cat_ FC) and ESMFold-predicted pLDDT. The no-penalty setting served as an unconstrained activity-maximization control, optimizing the kinetic predictor without any explicit foldability signal.

All three settings raised predicted activity above the wild-type baseline (*k*_cat_ FC = 1.0), confirming that CatESO provides a usable differentiable objective for improving predicted turnover. The gain obtained without structural regularization, however, came at a clear cost to predicted foldability. Taking pLDDT = 70 as the threshold for confident structural predictions, 82.0% and 86.0% of the sequences optimized with the ReLU and soft-hinge penalties, respectively, fell in the high-activity, high-confidence region, against only 36.0% of the unpenalized designs; the remaining 64.0% of the latter moved into a high-activity but low-confidence regime. The optimization trajectories (Fig.2D) show how this gap opens during the search: the no-penalty objective reached the largest predicted *k*_cat_ gains but steadily drove the search out of the high-confidence region, whereas both penalties rose more gradually and plateaued while holding pLDDT in check, yielding terminal distributions (Fig.2E) that were shifted toward higher *k*_cat_ FC only for the unpenalized setting, at lower and more dispersed pLDDT.

Folding-stability estimates from the Rosetta energy function gave an independent structural readout of the same effect (Fig.2F). The unpenalized designs were far more destabilizing, with a median folding ΔΔ*G* of 2.22kcal/mol, against 0.36kcal/mol and 0.65kcal/mol for the ReLU- and soft-hinge-regularized designs: without constraints, high predicted activity comes at the cost of global fold stability. Binding ΔΔ*G* values, by contrast, were small and comparable across the three settings (medians 0.23, 0.24, and 0.28 kcal/mol), so the optimized substitutions did not substantially remodel the predicted substrate-binding interface. Structural regularization thus preserved global fold stability without blocking gains in predicted turnover.

To confirm that this effect is not an artifact of scoring designs with the same models used to optimize them, we re-evaluated the three settings with two held-out evaluators, Protenix^83^ for structural confidence and OmniESI for predicted activity (Fig. S1). The same separation held: under OmniESI, the unpenalized designs again concentrated in a high-activity regime, while both penalties stayed closer to the wild-type baseline. Although every Protenix-predicted pLDDT remained above the confidence threshold, the unpenalized distribution sat systematically lower than those of the two regularized settings, echoing the foldability gap seen under ESMFold.

Mapping the selected substitutions onto the Q9P9E3 structure explained how the penalties shaped the search (Fig. 2G). The two regularized settings concentrated mutations within a compact, largely shared set of positions, whereas the unpenalized setting explored a broader and more dispersed set of residues, including sites likely to matter for maintaining the fold. Together, these analyses show that CatESO can use a differentiable substrate-conditioned kinetic predictor to increase predicted turnover, but that structural constraints are needed to prevent adversarial drift toward unstable sequences. We therefore adopted the structurally regularized design protocol for the applications that follow.

### CatESO enriches active, foldable variants for out-of-distribution enzymes

We built a stringent out-of-distribution (OOD) benchmark from RealKcat to assess how well CatESO generalizes beyond its training distribution. Within each of the seven top-level EC classes, we clustered the OOD enzymes and selected the centroid of the largest cluster, giving seven representative wild-type enzymes that span EC 1–7 (Fig. 3A, see Data curation). To minimize target-specific memorization, we excluded both the query enzymes and their close homologues from training. For each enzyme we relaxed the mutable Gumbel–Softmax logits under two sampling schedules: a fixed temperature (*τ* = *τ*_0_) and an annealed schedule that progressively lowers *τ* to sharpen the residue-level categorical distributions (Fig.3B). The two schedules trade broad exploration against late-stage refinement. After removing redundant sequences, we applied a sequence-distance penalty to keep variants close to their wild type, since enzymes tolerate only limited sequence perturbation;^39,84–86^ most retained designs carried 1–10 substitutions.

**Figure 3:**
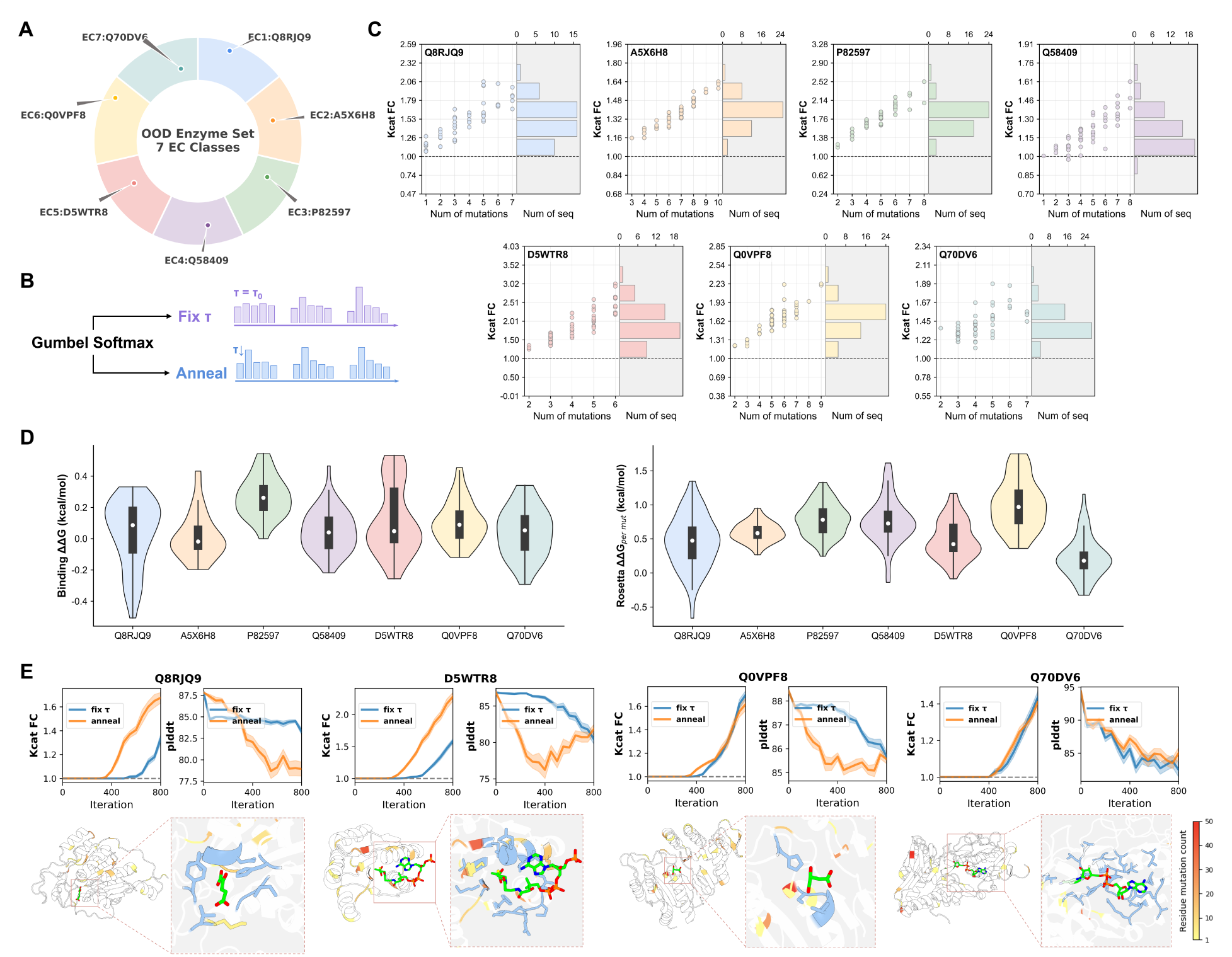
Out-of-distribution enzyme sequence optimization across the seven EC classes. (A) Construction of the OOD benchmark. Within each of the seven top-level EC classes, OOD enzymes were clustered and the centroid of the largest cluster was taken as the representative wild-type enzyme. (B) The two sampling schedules used to draw sequences from the Gumbel–Softmax logits. (C) CatESO-predicted *k*_cat_ fold changes for the non-redundant optimized designs of each enzyme, normalized to the corresponding wild type (grey dashed line, FC = 1.0). Each point relates a design to its number of substitutions; marginal histograms summarize design counts. (D) Energetic evaluation of the optimized designs for each enzyme. Violin plots show the binding ΔΔ*G* (left) and the per-mutation Rosetta ΔΔ*G* (right). (E) Per-enzyme optimization trajectories and structures for the representative targets. Top panels trace the CatESO-predicted *k*_cat_ fold change and the mean ESMFold pLDDT of the structurally relaxed sequences under the fixed *τ* and annealed schedules; solid lines denote the mean, shaded regions the ± s.d., and grey dashed lines the wild-type baseline. Bottom panels show the predicted structures of representative optimized variants.

Across the non-redundant designs, CatESO consistently enriched variants with higher predicted catalytic activity (Fig.3C). Overall, 98.8% of generated variants exceeded the predicted *k*_cat_ of their wild type, with a median predicted *k*_cat_ fold change of 1.52 and a maximum of 3.76 (Fig.S2A). The improvement held across all seven OOD enzymes, indicating that the enrichment reflects transferable sequence–activity relationships rather than target-specific effects.

To test whether these gains were specific to the CatESO scoring function or held under an independent model, we rescored all CatESO-generated sequences with OmniESI (Fig.S2B,S2C). OmniESI was more conservative but still showed an overall shift toward higher predicted activity, scoring five of the seven enzymes above their wild-type baseline. The clearest disagreement was Q58409, for which OmniESI assigned 96% of designs a predicted *k*_cat_ at or below wild type (Fig. S1B). Because both models were trained on the same underlying dataset and all seven test enzymes were held out from both training distributions, this discordance points to target-specific uncertainty in OOD extrapolation rather than memorization. It likely reflects differences in task formulation, label usage, or inductive bias—how each model scores mutations in sparsely supported regions of sequence space.

We next asked whether these activity gains came at a structural or binding cost, evaluating the folding penalty and binding-affinity change of the optimized designs for every OOD enzyme (Fig.3D). Because designs differ in the number of substitutions they carry, we report the Rosetta penalty per mutation, which isolates the average destabilization of a single substitution independent of design size. The per-mutation Rosetta ΔΔ*G* stayed mild across all seven enzymes, with medians between 0.18 and 0.97kcal/mol, so individual CatESO substitutions are on average close to thermoneutral even out of distribution. Binding ΔΔ*G* was likewise small, with medians near zero for every enzyme (between −0.02 and 0.26kcal/mol) and frequently negative minima: the substitutions left predicted substrate affinity essentially unchanged and occasionally improved it. Together, these per-enzyme energetics extend the single-enzyme behaviour of Fig.2 to the full OOD panel: CatESO raises predicted turnover while keeping individual substitutions fold-compatible and the substrate-binding interface intact.

To see how these designs emerge during optimization, we examined the trajectories of the representative enzymes (Fig.3E, Fig.S3). Under both schedules the predicted *k*_cat_ fold change rose progressively, but annealing climbed faster and produced mutant sequences at earlier steps, consistent with broad high-temperature exploration followed by low-temperature refinement. Mean pLDDT decreased modestly relative to the wild-type starting structures, as expected from the structural cost of sequence perturbation; the dip was larger under annealing, where early high-temperature steps sampled lower-confidence but higher-scoring regions, and pLDDT partially recovered as *τ* fell. Final variants nonetheless stayed within high- or acceptable-confidence structural regimes, and the accompanying structure views confirm that the accumulated substitutions left the overall fold intact (Fig. 3E).

We next examined where these substitutions fall on the enzyme structures. For each of the seven OOD enzymes, we mapped residue-level mutation frequencies from all designed variants onto the wild-type structure (Fig.3E, Fig.S3). The resulting patterns were sparse and highly non-uniform: mutations occurred at fewer than 30 residue positions per en-zyme—roughly 5–20% of the protein length—and clustered at a few recurrent hotspots that dominated the overall distribution, including V50 and T143 in A5X6H8, L96 in P82597, I190 in Q0VPF8, and G241 in Q70DV6 (Fig.S4). This sparsity shows that CatESO-derived optimization is driven by position-specific signals rather than broad or random mutational sampling.

These hotspots were not only sparse but structurally patterned: most fell outside the ligand-contacting catalytic core. This is consistent with the strong functional constraints typically imposed on catalytic residues and the local active-site geometry,^87,88^ where substitutions may disrupt substrate anchoring, catalytic chemistry, or transition-state stabilization.^89–91^ The relative conservation of the catalytic pocket suggests that CatESO tends to spare key catalytic determinants while still finding activity-enhancing substitutions. The enriched substitutions instead lay preferentially in residues adjacent to the binding pocket, in flexible loop regions near putative substrate-access channels, or in more distal folded regions (Fig. 4D). Substitutions at such sites can plausibly modulate activity without remodelling the core, acting through conformational or allosteric coupling^16,92^ that shifts substrate positioning, the population of catalytically competent conformations, or product release.^93,94^ These localization patterns should be read as predictive hypotheses, not direct evidence of catalytic effects: the structural analysis supports the plausibility of CatESO-selected substitutions, but the contribution of individual hotspots to *k*_cat_ will require experimental validation by site-directed mutagenesis and steady-state kinetics.

**Figure 4:**
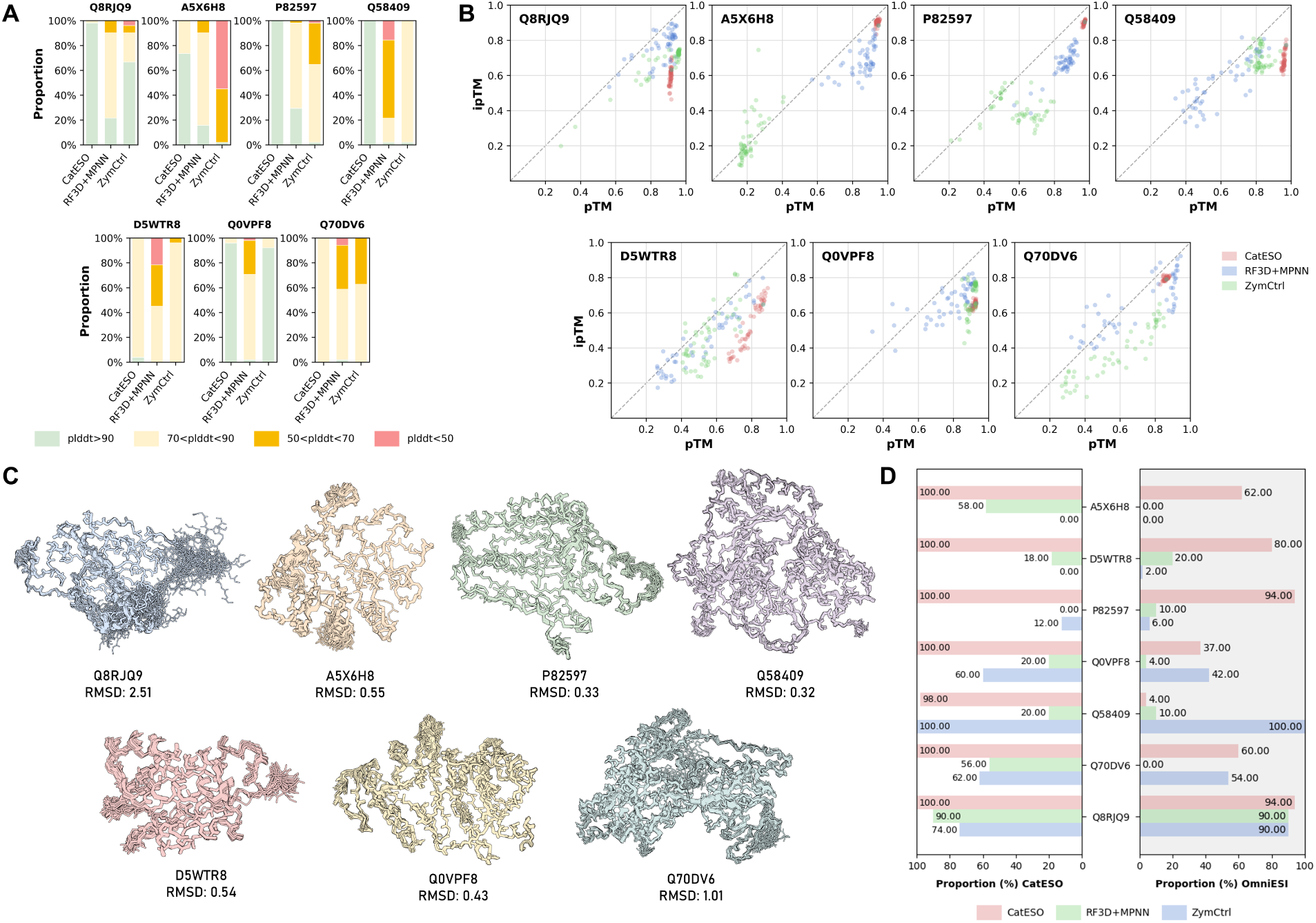
Structural and functional benchmarking of generated OOD enzyme variants. In all panels, non-redundant variants of seven OOD enzymes were generated by CatESO, RF3D+LigandMPNN, and ZymCtrl. (A) Distribution of pLDDT scores, shown as the proportion of variants within each pLDDT confidence interval. (B) Per-variant pTM and ipTM scores, each point denoting one variant, comparing global structural confidence (pTM) and predicted interface confidence (ipTM) across methods. (C) Backbone superposition of generated mutants onto the corresponding wild-type structure; main-chain traces were aligned and backbone RMSD computed per variant. (D) Proportion of variants that are both well-folded (pLDDT *>* 70) and predicted to be more active (*k*_cat_ fold change *>* 1.0), with fold change scored by CatESO (left) and OmniESI (right).

### CatESO outperforms structure- and language-based generators in joint activity–structure evaluation

We next benchmarked CatESO-designed variants against two baseline generators—RFdiffusion3^95^ coupled with LigandMPNN (RF3D+MPNN), and ZymCtrl—using Protenix^83^ to assess predicted structural plausibility across the seven OOD enzymes. Structural confidence varied markedly across the three methods (Fig.4A). Every CatESO variant exceeded pLDDT 70, and 66.9% reached the high-confidence regime above pLDDT 90, with little variation between enzymes. RF3D+MPNN and ZymCtrl were less consistent: only 9.1% and 22.6% of their designs exceeded pLDDT 90, while 32.8% and 26.9% fell below pLDDT 70 (Fig.S5A, S5B). Both baselines produced structurally plausible sequences for some targets, but only CatESO maintained high predicted confidence across all seven.

The same ranking held at the global and interface level (Fig. 4B). CatESO ranked highest, with median pTM and ipTM of 0.92 and 0.69 and 91.9% of designs above pTM 0.8. RF3D+MPNN was intermediate (median pTM/ipTM 0.82/0.66; 55.1% above 0.8), and ZymCtrl lowest (0.72/0.52; 39.4% above 0.8). CatESO thus improves predicted catalytic activity without sacrificing fold- or interface-level confidence, and does so more consistently than either baseline.

To determine whether CatESO’s high-confidence predictions reflect genuine preservation of the native tertiary fold rather than confident but non-native structures, we superposed the Protenix-predicted backbones of CatESO variants onto their wild-type structures (Fig.4C). Six of seven targets matched their wild-type folds closely, with backbone RMSD below 1.1Å: A5X6H8 (0.55 Å), P82597 (0.33 Å), Q58409 (0.32 Å), D5WTR8 (0.54 Å), Q0VPF8 (0.43 Å), and Q70DV6 (1.01 Å). The designs retained the native backbone across diverse scaffolds despite extensive sequence divergence. Q8RJQ9 was the only clear outlier (RMSD = 2.51 Å), which may indicate greater conformational flexibility or weaker structural constraints for this target. Together with the pLDDT, pTM, and ipTM results above, these superpositions show that CatESO optimization preserves native-like tertiary structure while improving predicted activity.

To combine the activity and structural criteria, we measured the fraction of variants that both passed a structural-confidence threshold and improved predicted *k*_cat_ over wild type (Fig.4D). Scored by CatESO, CatESO-optimized variants succeeded at a mean rate of 99.7% across the seven enzymes, compared with 37.4% for RF3D+MPNN and 44.0% for ZymCtrl (Fig.S3C, S3B), and led on six of seven targets. Because CatESO scores its own designs, we repeated the joint analysis with OmniESI as an independent activity evaluator. Absolute success rates dropped under this more conservative rescoring, but the ranking held. CatESO kept a positive median fold change, with 61.4% of structurally filtered variants above the wild-type baseline (Fig.S5C). RF3D+MPNN, which had been favoured under CatESO scoring for some targets, fell to a median OmniESI fold change of 0.48, with only 19.1% of designs above baseline. Much of its apparent gain therefore reflects CatESO-specific extrapolation rather than predictor-transferable improvement. ZymCtrl sat near baseline. CatESO again achieved the highest mean joint success rate, 61.4% versus 19.1% for RF3D+MPNN and 42.0% for ZymCtrl, and remained best on five of seven enzymes (Fig.S3C).

The per-enzyme breakdown shows how much uncertainty remains in model-based extrapolation (Fig.4D). Q58409 was the clearest predictor-dependent case: within the structurally filtered subset, CatESO assigned a median fold change of 1.20, which OmniESI reduced to 0.94. A variant judged structurally plausible and activity-enhancing by one model can thus be scored as neutral or deleterious by an independently trained one. The value of CatESO is therefore not the highest nominal predicted activity, but variants that simultaneously preserve structural plausibility, improve predicted activity, and retain support from an independent activity model across most OOD enzymes.

### Ablation study on key components

To quantify the contribution of each component in the CatESO feature-fusion module, we ran component-wise ablations. We removed MHSA, MCDC and ICFE either individually or together (N/ALL), holding all other architectural choices, optimization settings and evaluation protocols fixed. Each ablated model was evaluated under both the simple-fusion (SF) and concatenation-fusion (CF) heads, across the in-distribution split (100% sequence identity) and four increasingly OOD splits defined by maximum sequence identity to the training set. We compared each against the corresponding full SF/CatESO and CF/CatESO models, and against their ensemble, CatESO.

The three fusion components contribute mainly to generalization under distribution shift, not to in-distribution fitting (Table 1). On the ID split, ablated and full models performed similarly, with *R*^2^ clustered around 0.62–0.64 and PCC around 0.79–0.80. The gap widened as the sequence-identity threshold tightened from 99% to 40%, so the benefit of MHSA, MCDC and ICFE is clearest on more remote enzyme sequences. Removing all three modules at once gave the weakest OOD performance under both fusion heads: at the most stringent OOD-40% split, *R*^2^ fell to 0.287 for CF and 0.289 for SF, against 0.338 for the full CatESO ensemble. Among the single-component ablations, dropping ICFE cost the most under the strictest setting, with OOD-40% *R*^2^ of 0.316 for CF and 0.310 for SF. Explicit bidirectional modelling of enzyme–substrate interactions thus matters most when extrapolating to distant enzyme sequences. Removing MHSA or MCDC degraded performance more moderately but just as consistently, in line with their roles in contextualizing multi-molecule substrate representations and in substrate-conditioned modelling of the enzyme branch.

**Table 1:** Ablation analysis of CatESO for ID and OOD *k*_cat_ prediction. The upper table reports PCC and MAE, and the lower table reports RMSE and *R*^2^. Test splits were defined by the maximum sequence identity to the training set, with 100% used for ID evaluation and 99%, 80%, 60% and 40% for progressively stricter OOD evaluation. CF denotes concat fusion, SF denotes simple fusion, and N indicates that the corresponding component was removed.

| Model | ID - 100% |  | OOD - 99% |  | OOD - 80% |  | OOD - 60% |  | OOD - 40% |  |
| --- | --- | --- | --- | --- | --- | --- | --- | --- | --- | --- |
| | PCC ↑ | MAE ↓ | PCC ↑ | MAE ↓ | PCC ↑ | MAE ↓ | PCC ↑ | MAE ↓ | PCC ↑ | MAE ↓ |
| CF/N/MCDC | 0.798±0.001 | 0.674±0.001 | 0.645±0.004 | 0.971±0.002 | 0.642±0.004 | 1.000±0.002 | 0.614±0.007 | 1.087±0.002 | 0.583±0.007 | 1.125±0.001 |
| CF/N/MHSA | 0.799±0.000 | 0.671±0.002 | 0.646±0.003 | 0.963±0.005 | 0.645±0.004 | 0.991±0.005 | 0.615±0.005 | 1.089±0.006 | 0.585±0.004 | 1.131±0.007 |
| CF/N/ICFE | 0.796±0.001 | 0.679±0.002 | 0.641±0.003 | 0.973±0.004 | 0.639±0.004 | 0.999±0.004 | 0.610±0.004 | 1.089±0.005 | 0.577±0.008 | 1.129±0.003 |
| CF/N/ALL | 0.790±0.001 | 0.690±0.001 | 0.629±0.002 | 0.997±0.004 | 0.621±0.002 | 1.025±0.004 | 0.591±0.002 | 1.120±0.005 | 0.553±0.005 | 1.155±0.007 |
| SF/N/MCDC | 0.798±0.001 | 0.678±0.001 | 0.649±0.003 | 0.969±0.004 | 0.645±0.003 | 0.996±0.002 | 0.620±0.005 | 1.083±0.003 | 0.598±0.007 | 1.114±0.007 |
| SF/N/MHSA | 0.798±0.001 | 0.673±0.002 | 0.644±0.000 | 0.972±0.002 | 0.642±0.002 | 1.002±0.003 | 0.611±0.004 | 1.102±0.007 | 0.585±0.007 | 1.141±0.009 |
| SF/N/ICFE | 0.795±0.001 | 0.681±0.002 | 0.640±0.003 | 0.979±0.004 | 0.638±0.005 | 1.006±0.004 | 0.609±0.006 | 1.104±0.007 | 0.575±0.010 | 1.141±0.009 |
| SF/N/ALL | 0.789±0.002 | 0.693±0.002 | 0.628±0.003 | 0.995±0.004 | 0.620±0.003 | 1.021±0.004 | 0.588±0.004 | 1.119±0.007 | 0.554±0.007 | 1.153±0.005 |
| CF/CatESO | 0.798±0.001 | 0.672±0.000 | 0.648±0.005 | 0.966±0.007 | 0.647±0.005 | 0.992±0.008 | 0.618±0.006 | 1.088±0.009 | 0.588±0.008 | 1.133±0.012 |
| SF/CatESO | 0.799±0.000 | 0.671±0.002 | 0.649±0.003 | 0.956±0.004 | 0.649±0.004 | 0.982±0.005 | 0.621±0.006 | 1.078±0.005 | 0.595±0.009 | 1.121±0.010 |
| CatESO | 0.799±0.000 | 0.670±0.001 | 0.650±0.003 | 0.959±0.004 | 0.650±0.005 | 0.984±0.007 | 0.623±0.008 | 1.077±0.009 | 0.601±0.009 | 1.116±0.009 |
| | RMSE ↓ | $R^2$ ↑ | RMSE ↓ | $R^2$ ↑ | RMSE ↓ | $R^2$ ↑ | RMSE ↓ | $R^2$ ↑ | RMSE ↓ | $R^2$ ↑ |
| CF/N/MCDC | 0.985±0.003 | 0.634±0.002 | 1.331±0.006 | 0.406±0.006 | 1.340±0.006 | 0.402±0.005 | 1.425±0.008 | 0.365±0.007 | 1.449±0.007 | 0.324±0.007 |
| CF/N/MHSA | 0.984±0.001 | 0.635±0.001 | 1.322±0.004 | 0.413±0.003 | 1.331±0.004 | 0.410±0.004 | 1.424±0.005 | 0.366±0.004 | 1.448±0.003 | 0.325±0.003 |
| CF/N/ICFE | 0.991±0.003 | 0.630±0.002 | 1.332±0.004 | 0.405±0.003 | 1.339±0.004 | 0.403±0.004 | 1.428±0.005 | 0.363±0.004 | 1.457±0.007 | 0.316±0.007 |
| CF/N/ALL | 1.005±0.002 | 0.620±0.001 | 1.363±0.003 | 0.377±0.002 | 1.375±0.003 | 0.371±0.002 | 1.458±0.002 | 0.336±0.002 | 1.488±0.006 | 0.287±0.005 |
| SF/N/MCDC | 0.986±0.003 | 0.634±0.002 | 1.323±0.005 | 0.413±0.005 | 1.333±0.004 | 0.408±0.004 | 1.416±0.008 | 0.374±0.007 | 1.428±0.008 | 0.343±0.008 |
| SF/N/MHSA | 0.985±0.003 | 0.634±0.002 | 1.326±0.001 | 0.410±0.001 | 1.337±0.002 | 0.405±0.001 | 1.433±0.005 | 0.359±0.004 | 1.453±0.008 | 0.320±0.007 |
| SF/N/ICFE | 0.992±0.002 | 0.629±0.002 | 1.334±0.004 | 0.402±0.004 | 1.342±0.006 | 0.400±0.006 | 1.435±0.007 | 0.357±0.006 | 1.464±0.010 | 0.310±0.009 |
| SF/N/ALL | 1.007±0.003 | 0.618±0.002 | 1.364±0.003 | 0.375±0.003 | 1.376±0.003 | 0.369±0.003 | 1.463±0.005 | 0.331±0.004 | 1.486±0.007 | 0.289±0.006 |
| CF/CatESO | 0.986±0.002 | 0.633±0.002 | 1.320±0.008 | 0.415±0.007 | 1.328±0.008 | 0.413±0.007 | 1.421±0.009 | 0.370±0.008 | 1.444±0.012 | 0.328±0.011 |
| SF/CatESO | 0.984±0.001 | 0.635±0.001 | 1.318±0.004 | 0.417±0.004 | 1.326±0.004 | 0.415±0.004 | 1.418±0.005 | 0.372±0.004 | 1.440±0.007 | 0.333±0.007 |
| CatESO | 0.983±0.001 | 0.636±0.001 | 1.318±0.005 | 0.417±0.004 | 1.326±0.008 | 0.415±0.007 | 1.416±0.010 | 0.374±0.009 | 1.434±0.009 | 0.338±0.009 |

We further evaluated whether combining the two fusion heads could enhance predictive robustness. The final CatESO model ensembles the SF/CatESO and CF/CatESO predictors, and performed best overall. It reached the highest PCC on every OOD split—0.650, 0.650, 0.623 and 0.601 at the 99%, 80%, 60% and 40% thresholds—and the best ID performance on all four metrics (PCC 0.799, MAE 0.670, RMSE 0.983, *R*^2^ 0.636). On the OOD splits, the ensemble was best or tied-best in *R*^2^ and RMSE at 99%, 80% and 60%, and beat both single-head models on every metric at the strictest 40% split, where its *R*^2^ of 0.338 exceeded 0.333 for SF/CatESO and 0.328 for CF/CatESO. Combining the two complementary sequence–substrate fusion representations therefore pays off precisely where extrapolation is hardest.

These ablations point to three complementary inductive biases in CatESO. ICFE-mediated interaction modelling is the single largest contributor to OOD generalization; MHSA and MCDC add substrate-contextual and substrate-conditioned enzyme representations on top of it. Ensembling the two fusion heads buys a further margin of robustness, and gives the most consistent performance as the OOD splits grow stricter.

## Methods

### Dataset curation

#### Training data

We used CatPred-DB as our source of training data, taking the curated *k*_cat_ dataset together with the original train/validation/test splits released with CatPred. CatPred-DB was assembled from experimentally reported kinetic measurements in BRENDA (release 2022.2) and SABIO-RK records through November 2023. It covers wild-type enzymes with mapped protein sequences and canonicalized reactant SMILES; the processed *k*_cat_ subset contains 23,197 curated in vitro measurements.

#### Stratified evaluation

To probe how predictive accuracy varies across enzyme families and sequence lengths, we evaluated CatESO on stratified subsets of the CatPred-DB test set. We grouped enzymes by family using the first digit of the EC number, yielding EC1–EC5 and a merged EC6–7 class (EC7 alone held too few test entries to evaluate reliably). Independently, we binned enzymes by sequence length into short (*<* 300 residues), medium (300–700) and long (*>* 700). Both stratifications inherit the ID and OOD splits of the full test set.

#### Strict out-of-distribution benchmark

To test whether CatESO generalizes beyond the CatPred training distribution, we built a strict OOD benchmark from RealKcat. We first discarded any RealKcat entry that matched a CatPred sequence exactly or shared a UniProt accession with the CatPred training, validation, or test splits, then computed ESMFold pLDDT scores for the survivors. From these we kept wild-type enzymes meeting four criteria: a valid UniProt identifier, log_10_(*k*_cat_) within [−1.5, 1.5], a sequence of 100–700 residues, and a wild-type ESMFold pLDDT of at least 70. This left 582 candidates.

To suppress residual remote homology, we aligned these candidates against the pooled CatPred sequences (train, validation, and test) with MMseqs2 easy-search, removing any candidate whose best hit reached both ≥ 40% sequence identity and ≥ 80% bidirectional coverage. The resulting strict-OOD benchmark holds 269 wild-type enzymes spanning all seven primary EC classes.

#### Optimization scaffolds

For the sequence-optimization experiments, we built a compact, class-balanced benchmark by clustering the strict-OOD enzymes within each primary EC class (MMseqs2, 40% identity, 80% bidirectional coverage) and drawing one representative per class—seven starting scaffolds in total. Q9P9E3, also from the strict-OOD set, served separately as a moderate-confidence scaffold for the illustrative optimization example.

### Overview of the Proposed Framework

CatESO was designed to integrate enzyme sequence information and substrate molecular structure for catalytic-turnover prediction and differentiable enzyme sequence optimization. For kinetic prediction, enzyme sequences were encoded with a frozen ESM-2 (3B) model to obtain residue-level protein representations.^82,96^ In parallel, each substrate, co-substrate or reactant molecule was represented as a molecular graph and processed by a residual graph convolutional network (GCN) to generate atom-level features, which were subsequently pooled into molecule-level embeddings.^97^ For reactions containing multiple molecular components, multi-head self-attention (MHSA)^98^ was applied across molecule-level embeddings to model dependencies within the reactant set and to generate contextualized molecular representations.

Enzyme and reactant representations were integrated using two cross-modal modules. First, a molecule-conditioned dynamic convolution module (MCDC) used contextualized reactant embeddings to parameterize convolutional filters over residue-level protein features, enabling substrate-dependent extraction of local sequence patterns. Second, an interaction-cognition feature extraction module (ICFE) refined enzyme and molecular representations through bidirectional cross-attention, allowing protein residues and molecular features to attend to each other. The resulting interaction fingerprint, which summarizes enzyme sequence context, reactant-set context and their learned compatibility, was passed to a multilayer prediction head to regress log_10_(*k*_cat_). To improve prediction robustness, multiple CatESO predictors with distinct hyperparameter configurations were trained, and their ensemble mean was used as the final estimate of catalytic turnover.

For enzyme design, the trained kinetic predictor was used as a fixed differentiable surrogate objective. During optimization, the kinetic predictor, ESM-2 encoder and, when used, the ESMFold-based structural evaluator were kept frozen, whereas only the sequence logits at predefined mutable residue positions were updated. Regularization terms were optionally added to penalize excessive sequence deviation from the reference enzyme and low predicted structural confidence. This formulation allowed catalytic activity, sequence plausibility and structural feasibility to be optimized jointly within a single differentiable framework.

### Multi-Head Self-Attention Feature Modulation

Before cross-modal interaction, a multi-head self-attention block is applied to the substrate representation. This operation allows atom-level features to incorporate information from non-adjacent atoms in the molecular graph, thereby complementing the local structural information encoded by graph convolution. The resulting substrate representation provides the molecular context used by subsequent enzyme feature extraction modules.

### Molecular-Guided Dynamic Convolution

The molecular–guided dynamic convolution module updates the enzyme representation through substrate–conditioned convolution along the protein sequence. Rather than using fixed convolutional kernels shared across all enzyme–substrate pairs, the module generates depthwise convolution kernels from a pooled substrate descriptor. In this manner, the convolution operation is conditioned on the molecular context of the substrate. For each convolutional scale, the substrate descriptor is mapped to a set of dynamic depthwise kernels and applied to the enzyme sequence representation. Outputs from multiple kernel sizes are aggregated and normalized to obtain the updated enzyme features. This module provides a mechanism for modelling substrate-dependent local residue patterns before subsequent cross-modal feature enhancement (Fig. 2B).

### Interactive Cross-Modal Feature Enhancement

After modality-specific refinement, ICFE module is used to model bidirectional interactions between substrate atoms and enzyme residues. Each block consists of cross-attention in both directions, followed by modality-specific feed-forward refinement (Fig. 2C).

To combine intra-modal and cross-modal information, the module uses learnable scalar coefficients rather than fixed residual addition:

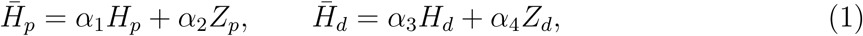

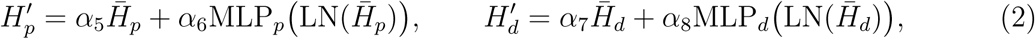

where *Z_p_* and *Z_d_* denote the cross-attended enzyme and substrate features, respectively, and *α*_1_*, . . ., α*_8_ are learnable coefficients. Multiple ICFE blocks are stacked to progressively refine the cross-modal representations.

### Prediction Head

The refined substrate and enzyme representations are converted into fixed-length vectors using masked average pooling over valid tokens. The resulting modality-level features, *H_d_* and *H_p_*, are combined by either element-wise addition or feature concatenation and then passed to the prediction head for catalytic turnover estimation.

### Constrained Differentiable Sequence Optimization

For enzyme sequence design, the trained predictor is used as a fixed differentiable surrogate objective. During optimization, the *k_cat_* predictor, ESM-2 encoder and optional ESMFold-based evaluator are kept frozen, and only the sequence variables at predefined mutable positions are updated.

The search space is restricted to the 20 canonical amino acids. For each mutable residue position, a trainable logit vector is defined over this amino-acid set, yielding a logit matrix Θ_seq_ ∈ ℝ*^N^*^mut^*^×^*^20^, where *N*_mut_ is the number of mutable positions. Fixed residues are clamped to their original identities throughout optimization, allowing prior residue-level constraints to be incorporated directly into the parameterization.

At each optimization step, the amino-acid logits are mapped to the ESM-2 token space, while non-canonical residues and special tokens are masked with large negative logits. A differentiable sequence representation is then sampled using the Gumbel–Softmax relaxation:

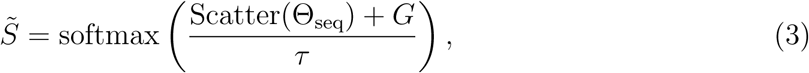

where *G* denotes Gumbel noise and *τ* is the temperature parameter. This relaxation permits gradient-based updates while retaining a categorical sequence parameterization.

The relaxed sequence representation is projected into the ESM-2 embedding space and encoded by the frozen ESM-2 model. The resulting contextualized enzyme features, together with the fixed substrate graph representation, are passed to the frozen predictor to compute the optimization objective under the specified residue constraints.

### Sequence and Structural Regularization

CatESO formulates enzyme sequence optimization as a regularized objective that maximizes the predicted catalytic activity while optionally constraining sequence divergence and structural confidence. The catalytic objective is defined as the negative predicted activity:

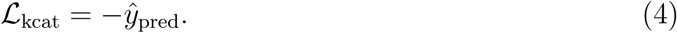

To limit the mutational burden and preserve similarity to the starting enzyme, we included an optional sequence regularization term that penalizes deviations from the initial sequence:

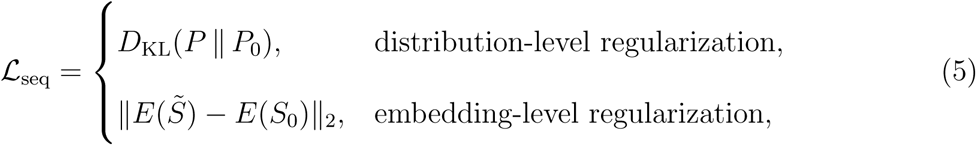

where *P* and *P*_0_ denote the optimized and initial residue distributions, respectively, *S̃* is the optimized soft sequence, *S*_0_ is the starting sequence, and *E*(·) is the frozen ESM-2 encoder. This term restricts the optimized sequence to remain close to the wild-type scaffold, thereby reducing excessive sequence drift and limiting the number of introduced substitutions.

When structural regularization is enabled, a frozen ESMFold-based module is used to estimate a structural confidence score *p*, such as pLDDT. Structural degradation is penalized through a threshold-based loss:

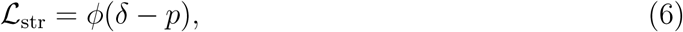

where *δ* is the confidence threshold and *ϕ* is selected from ReLU, softplus or soft hinge. These choices trade off computational efficiency and smoothness: ReLU is inexpensive but non-smooth at the threshold, whereas softplus provides a smooth relaxation with higher computational cost. Soft hinge serves as an intermediate option, smoothing the transition near *δ* while being clamped to zero, with zero gradient, once *p > δ* + *m*. This avoids unnecessary back-propagation through the ESMFold branch for sequences that already satisfy the structural confidence constraint.

The final optimization objective is

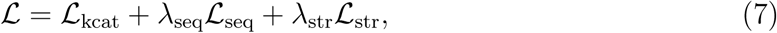

where *⋋*_seq_ and *⋋*_str_ control the strength of the sequence and structural regularization terms, respectively. This objective enables substrate-aware catalytic optimization while allowing explicit control over sequence conservation and structural plausibility.

### Optimization Procedure

During optimization, all parameters of kinetic predictor, ESM-2 and ESMFold, ^82^ are kept fixed, and only the sequence logits at mutable residue positions are updated. Each mutable position is parameterized by a logit vector over the 20 canonical amino acids. At each iteration, these logits are transformed into a differentiable soft sequence using a temperature-controlled Gumbel–Softmax relaxation.^77,79^ The relaxed amino-acid representation is projected into the ESM-2 embedding space and propagated through the frozen ESM-2 encoder. The resulting protein representation is combined with substrate molecular features and passed to the fixed catalytic-turnover predictor to compute the ensemble-predicted log_10_(*k*_cat_) objective.

The optimization loss includes the negative catalytic objective together with optional regularization terms that constrain sequence divergence from the reference enzyme and penalize low predicted folding confidence. Structural confidence is evaluated using ESMFold-derived pLDDT scores, with penalties applied when predicted folding quality falls below predefined thresholds. User-specified positions, including catalytic residues or experimentally validated binding-site residues, can be masked from optimization by fixing their logits to the reference amino acids. Gradients are back-propagated only to the mutable sequence logits, which are updated using a first-order optimizer such as Adam or AdamW. Temperature annealing can be applied during optimization to gradually sharpen the residue distribution. After optimization, a discrete enzyme sequence is obtained by selecting the amino acid with the highest logit at each mutable position.

### Implementation

CatESO was implemented in Python 3.7 with PyTorch 1.12.1 and consists of two components: a substrate-conditioned *k*_cat_ predictor and a gradient-based enzyme sequence optimizer. In the predictor, enzyme sequences were encoded using the ESM-2-3B protein language model, whereas substrates were represented by graph convolutional networks implemented with DGL^99^ and DGL-LifeSci.^100^ The predictor was trained using the Adam optimizer with a cosine learning-rate schedule and data-parallel training on four NVIDIA A800 GPUs. During sequence optimization, both the trained kinetic predictor and the protein structure model were kept frozen. Predictor inference was performed on a single NVIDIA RTX 4090D GPU, whereas CatESO-based sequence optimization was conducted on a single NVIDIA A800 GPU. Representative protein structures were visualized using PyMOL.^101^

## Data and code availability

The enzyme kinetic parameter datasets used in this study are publicly available from previously published studies.^42,43^ The processed datasets, source code, trained models and scripts required to reproduce the main analyses are available on GitHub at https://github.com/zhenjiagan/CatESO.git.

## Acknowledgement

We gratefully acknowledge the high-performance computing resources provided by NYU Shanghai, NYU Abu Dhabi, and the NYU Torch supercomputing cluster at New York University, as well as the expert technical support provided by their dedicated staff.

## Supporting Information Available

**Table S1:** Comparison of CatESO with published baseline methods for ID and OOD *k_cat_* prediction on the CatPred-DB test set. The upper panel reports PCC and MAE, and the lower panel reports RMSE and *R*^2^; arrows indicate whether higher (↑) or lower (↓) values are better. Test splits were defined by the maximum sequence identity to the training set, with 100% used for ID evaluation and 99%, 80%, 60% and 40% for progressively stricter OOD evaluation. Baselines are DLKcat, UniKP, CatPred and OmniESI. Values are mean ± standard deviation across independent runs.

| Model | ID - 100% |  | OOD - 99% |  | OOD - 80% |  | OOD - 60% |  | OOD - 40% |  |
| --- | --- | --- | --- | --- | --- | --- | --- | --- | --- | --- |
| | PCC ↑ | MAE ↓ | PCC ↑ | MAE ↓ | PCC ↑ | MAE ↓ | PCC ↑ | MAE ↓ | PCC ↑ | MAE ↓ |
| DLKcat | 0.548±0.137 | 0.999±0.147 | 0.463±0.077 | 1.200±0.127 | 0.418±0.097 | 1.242±0.128 | 0.388±0.106 | 1.330±0.134 | 0.370±0.071 | 1.323±0.085 |
| UniKP | 0.761±0.001 | 0.757±0.001 | 0.612±0.005 | 1.053±0.005 | 0.602±0.008 | 1.096±0.002 | 0.576±0.018 | 1.186±0.005 | 0.525±0.018 | 1.202±0.004 |
| CatPred | 0.775±0.001 | 0.715±0.002 | 0.625±0.003 | 0.999±0.014 | 0.606±0.006 | 1.045±0.017 | 0.581±0.011 | 1.145±0.023 | 0.588±0.017 | 1.163±0.030 |
| OmniESI | 0.795±0.002 | 0.688±0.002 | 0.643±0.010 | 0.968±0.005 | 0.638±0.011 | 1.000±0.008 | 0.611±0.011 | 1.091±0.005 | 0.556±0.014 | 1.152±0.010 |
| CatESO | 0.799±0.001 | 0.670±0.001 | 0.650±0.004 | 0.959±0.005 | 0.650±0.006 | 0.984±0.009 | 0.623±0.009 | 1.077±0.011 | 0.601±0.011 | 1.116±0.011 |
| | RMSE ↓ | $R^2$ ↑ | RMSE ↓ | $R^2$ ↑ | RMSE ↓ | $R^2$ ↑ | RMSE ↓ | $R^2$ ↑ | RMSE ↓ | $R^2$ ↑ |
| DLKcat | 1.329±0.154 | 0.284±0.156 | 1.629±0.143 | 0.156±0.142 | 1.687±0.148 | 0.110±0.142 | 1.786±0.155 | 0.085±0.133 | 1.757±0.149 | 0.049±0.111 |
| UniKP | 1.067±0.002 | 0.571±0.001 | 1.394±0.005 | 0.347±0.004 | 1.432±0.005 | 0.317±0.005 | 1.526±0.011 | 0.273±0.011 | 1.557±0.012 | 0.219±0.012 |
| CatPred | 1.037±0.001 | 0.600±0.001 | 1.353±0.005 | 0.389±0.004 | 1.394±0.009 | 0.364±0.008 | 1.490±0.014 | 0.328±0.013 | 1.448±0.018 | 0.322±0.016 |
| OmniESI | 0.991±0.004 | 0.630±0.003 | 1.326±0.015 | 0.410±0.014 | 1.341±0.017 | 0.401±0.015 | 1.431±0.015 | 0.360±0.014 | 1.476±0.016 | 0.298±0.015 |
| CatESO | 0.983±0.001 | 0.636±0.001 | 1.318±0.006 | 0.417±0.005 | 1.326±0.009 | 0.415±0.008 | 1.416±0.012 | 0.374±0.010 | 1.434±0.011 | 0.338±0.011 |

**Table S2:** Per-EC-class comparison of CatESO with published baseline methods for ID and OOD *k_cat_* prediction on the CatPred-DB test set (EC1–EC3). Enzymes are grouped by the first digit of their EC number, and each group is evaluated separately. Within every group, PCC and *R*^2^ are reported; arrows indicate whether higher (↑) values are better. Test splits were defined by the maximum sequence identity to the training set, with 100% used for ID evaluation and 99%, 80%, 60% and 40% for progressively stricter OOD evaluation. Baselines are DLKcat, UniKP, CatPred and OmniESI. Values are mean ± standard deviation across independent runs. Continued in the next table.

| EC1 |  |  |  |  |  |  |  |  |  |  |
| --- | --- | --- | --- | --- | --- | --- | --- | --- | --- | --- |
| Model | ID - 100% |  | OOD - 99% |  | OOD - 80% |  | OOD - 60% |  | OOD - 40% |  |
| | PCC ↑ | $R^2$ ↑ | PCC ↑ | $R^2$ ↑ | PCC ↑ | $R^2$ ↑ | PCC ↑ | $R^2$ ↑ | PCC ↑ | $R^2$ ↑ |
| DLKcat | 0.572±0.161 | 0.317±0.199 | 0.450±0.082 | 0.125±0.184 | 0.412±0.114 | 0.102±0.162 | 0.393±0.102 | 0.081±0.108 | 0.395±0.012 | 0.030±0.025 |
| UniKP | 0.803±0.003 | 0.633±0.005 | 0.579±0.010 | 0.329±0.009 | 0.566±0.011 | 0.285±0.010 | 0.552±0.015 | 0.245±0.007 | 0.539±0.025 | 0.200±0.013 |
| CatPred | 0.809±0.002 | 0.648±0.004 | 0.675±0.023 | 0.435±0.038 | 0.646±0.028 | 0.385±0.040 | 0.624±0.057 | 0.353±0.069 | 0.693±0.040 | 0.425±0.036 |
| OmniESI | 0.838±0.002 | 0.701±0.003 | 0.681±0.013 | 0.454±0.019 | 0.688±0.015 | 0.470±0.022 | 0.716±0.015 | 0.496±0.023 | 0.702±0.008 | 0.454±0.012 |
| CatESO | 0.838±0.002 | 0.701±0.003 | 0.667±0.010 | 0.428±0.016 | 0.671±0.015 | 0.446±0.021 | 0.693±0.021 | 0.473±0.025 | 0.706±0.033 | 0.471±0.031 |

| EC2 |  |  |  |  |  |  |  |  |  |  |
| --- | --- | --- | --- | --- | --- | --- | --- | --- | --- | --- |
| Model | ID - 100% |  | OOD - 99% |  | OOD - 80% |  | OOD - 60% |  | OOD - 40% |  |
| | PCC ↑ | $R^2$ ↑ | PCC ↑ | $R^2$ ↑ | PCC ↑ | $R^2$ ↑ | PCC ↑ | $R^2$ ↑ | PCC ↑ | $R^2$ ↑ |
| DLKcat | 0.459±0.174 | 0.161±0.203 | 0.323±0.089 | 0.012±0.117 | 0.292±0.090 | -0.012±0.100 | 0.262±0.091 | -0.022±0.100 | 0.180±0.102 | -0.131±0.148 |
| UniKP | 0.736±0.006 | 0.525±0.008 | 0.592±0.018 | 0.316±0.017 | 0.605±0.029 | 0.297±0.022 | 0.595±0.034 | 0.259±0.024 | 0.496±0.050 | 0.186±0.027 |
| CatPred | 0.725±0.004 | 0.522±0.006 | 0.482±0.009 | 0.225±0.015 | 0.508±0.015 | 0.246±0.018 | 0.505±0.027 | 0.234±0.019 | 0.437±0.052 | 0.160±0.042 |
| OmniESI | 0.742±0.005 | 0.545±0.008 | 0.517±0.015 | 0.250±0.020 | 0.554±0.017 | 0.289±0.021 | 0.528±0.017 | 0.262±0.018 | 0.393±0.029 | 0.113±0.027 |
| CatESO | 0.743±0.003 | 0.541±0.005 | 0.538±0.004 | 0.275±0.007 | 0.594±0.006 | 0.334±0.010 | 0.574±0.006 | 0.305±0.008 | 0.491±0.004 | 0.200±0.004 |

| EC3 |  |  |  |  |  |  |  |  |  |  |
| --- | --- | --- | --- | --- | --- | --- | --- | --- | --- | --- |
| Model | ID - 100% |  | OOD - 99% |  | OOD - 80% |  | OOD - 60% |  | OOD - 40% |  |
| | PCC ↑ | $R^2$ ↑ | PCC ↑ | $R^2$ ↑ | PCC ↑ | $R^2$ ↑ | PCC ↑ | $R^2$ ↑ | PCC ↑ | $R^2$ ↑ |
| DLKcat | 0.519±0.137 | 0.258±0.139 | 0.291±0.022 | -0.050±0.103 | 0.267±0.021 | -0.052±0.084 | 0.260±0.054 | -0.108±0.126 | 0.316±0.064 | -0.079±0.125 |
| UniKP | 0.731±0.003 | 0.525±0.004 | 0.586±0.017 | 0.234±0.008 | 0.529±0.026 | 0.161±0.011 | 0.515±0.043 | 0.124±0.023 | 0.438±0.060 | 0.098±0.029 |
| CatPred | 0.738±0.001 | 0.540±0.002 | 0.522±0.037 | 0.271±0.038 | 0.527±0.044 | 0.264±0.039 | 0.495±0.052 | 0.218±0.049 | 0.476±0.069 | 0.210±0.060 |
| OmniESI | 0.760±0.004 | 0.577±0.006 | 0.552±0.019 | 0.246±0.033 | 0.498±0.022 | 0.186±0.037 | 0.430±0.028 | 0.111±0.040 | 0.401±0.034 | 0.125±0.034 |
| CatESO | 0.781±0.001 | 0.609±0.003 | 0.604±0.008 | 0.309±0.005 | 0.561±0.006 | 0.252±0.010 | 0.505±0.008 | 0.187±0.014 | 0.485±0.021 | 0.186±0.020 |

| EC4 |  |  |  |  |  |  |  |  |  |  |
| --- | --- | --- | --- | --- | --- | --- | --- | --- | --- | --- |
| Model | ID - 100% |  | OOD - 99% |  | OOD - 80% |  | OOD - 60% |  | OOD - 40% |  |
| | PCC ↑ | $R^2$ ↑ | PCC ↑ | $R^2$ ↑ | PCC ↑ | $R^2$ ↑ | PCC ↑ | $R^2$ ↑ | PCC ↑ | $R^2$ ↑ |
| DLKcat | 0.683±0.035 | 0.440±0.010 | 0.710±0.136 | 0.499±0.212 | 0.711±0.161 | 0.487±0.271 | 0.697±0.200 | 0.407±0.372 | 0.701±0.121 | 0.405±0.256 |
| UniKP | 0.690±0.009 | 0.451±0.013 | 0.690±0.026 | 0.412±0.027 | 0.681±0.031 | 0.364±0.031 | 0.600±0.038 | 0.239±0.041 | 0.626±0.086 | 0.169±0.094 |
| CatPred | 0.824±0.004 | 0.675±0.007 | 0.757±0.009 | 0.570±0.013 | 0.790±0.002 | 0.613±0.002 | 0.822±0.004 | 0.602±0.012 | 0.888±0.027 | 0.680±0.051 |
| OmniESI | 0.797±0.003 | 0.624±0.005 | 0.776±0.015 | 0.593±0.025 | 0.799±0.012 | 0.611±0.026 | 0.792±0.017 | 0.500±0.037 | 0.828±0.023 | 0.476±0.059 |
| CatESO | 0.790±0.005 | 0.614±0.008 | 0.768±0.004 | 0.581±0.007 | 0.798±0.003 | 0.614±0.009 | 0.780±0.009 | 0.493±0.017 | 0.890±0.003 | 0.541±0.029 |

| EC5 |  |  |  |  |  |  |  |  |  |  |
| --- | --- | --- | --- | --- | --- | --- | --- | --- | --- | --- |
| Model | ID - 100% |  | OOD - 99% |  | OOD - 80% |  | OOD - 60% |  | OOD - 40% |  |
| | PCC ↑ | $R^2$ ↑ | PCC ↑ | $R^2$ ↑ | PCC ↑ | $R^2$ ↑ | PCC ↑ | $R^2$ ↑ | PCC ↑ | $R^2$ ↑ |
| DLKcat | 0.582±0.049 | 0.317±0.039 | 0.727±0.025 | 0.487±0.057 | 0.651±0.064 | 0.169±0.180 | 0.690±0.035 | 0.293±0.180 | 0.869±0.046 | 0.112±0.351 |
| UniKP | 0.665±0.022 | 0.354±0.025 | 0.734±0.025 | 0.402±0.025 | 0.613±0.045 | 0.257±0.056 | 0.637±0.047 | 0.341±0.050 | 0.693±0.108 | 0.117±0.104 |
| CatPred | 0.738±0.010 | 0.542±0.014 | 0.861±0.005 | 0.723±0.017 | 0.775±0.016 | 0.576±0.021 | 0.825±0.013 | 0.636±0.031 | 0.939±0.016 | 0.864±0.036 |
| OmniESI | 0.764±0.007 | 0.564±0.012 | 0.869±0.018 | 0.737±0.031 | 0.774±0.033 | 0.574±0.050 | 0.785±0.034 | 0.602±0.047 | 0.841±0.040 | 0.549±0.099 |
| CatESO | 0.790±0.000 | 0.602±0.003 | 0.845±0.004 | 0.692±0.001 | 0.710±0.007 | 0.469±0.004 | 0.713±0.013 | 0.496±0.015 | 0.696±0.008 | 0.368±0.037 |

| EC6-7 |  |  |  |  |  |  |  |  |  |  |
| --- | --- | --- | --- | --- | --- | --- | --- | --- | --- | --- |
| Model | ID - 100% |  | OOD - 99% |  | OOD - 80% |  | OOD - 60% |  | OOD - 40% |  |
| | PCC ↑ | $R^2$ ↑ | PCC ↑ | $R^2$ ↑ | PCC ↑ | $R^2$ ↑ | PCC ↑ | $R^2$ ↑ | PCC ↑ | $R^2$ ↑ |
| DLKcat | 0.459±0.161 | 0.111±0.173 | 0.603±0.327 | 0.203±0.670 | 0.603±0.327 | 0.203±0.670 | 0.424±0.632 | 0.286±0.492 | 0.601±0.404 | 0.183±0.586 |
| UniKP | 0.725±0.026 | 0.482±0.031 | 0.607±0.107 | 0.269±0.057 | 0.607±0.107 | 0.269±0.057 | 0.636±0.092 | 0.290±0.053 | 0.725±0.102 | 0.286±0.067 |
| CatPred | 0.716±0.024 | 0.481±0.030 | 0.555±0.040 | 0.300±0.037 | 0.555±0.040 | 0.300±0.037 | 0.487±0.050 | 0.219±0.042 | 0.454±0.061 | 0.029±0.068 |
| OmniESI | 0.785±0.010 | 0.612±0.013 | 0.696±0.025 | 0.462±0.031 | 0.696±0.025 | 0.462±0.031 | 0.658±0.031 | 0.404±0.034 | 0.582±0.056 | 0.259±0.052 |
| CatESO | 0.761±0.016 | 0.575±0.028 | 0.614±0.070 | 0.369±0.077 | 0.614±0.070 | 0.369±0.077 | 0.552±0.086 | 0.299±0.088 | 0.513±0.095 | 0.174±0.105 |

**Table S3:** Comparison of CatESO with published baseline methods for ID and OOD *k_cat_* prediction on the CatPred-DB test set, stratified by enzyme sequence length. Enzymes are grouped by amino-acid sequence length into short, medium and long classes, and each group is evaluated separately. Within every group, PCC and *R*^2^ are reported; arrows indicate whether higher (↑) values are better. Test splits were defined by the maximum sequence identity to the training set, with 100% used for ID evaluation and 99%, 80%, 60% and 40% for progressively stricter OOD evaluation. Baselines are DLKcat, UniKP, CatPred and OmniESI. Values are mean ± standard deviation across independent runs.

| Short |  |  |  |  |  |  |  |  |  |  |
| --- | --- | --- | --- | --- | --- | --- | --- | --- | --- | --- |
| Model | ID - 100% |  | OOD - 99% |  | OOD - 80% |  | OOD - 60% |  | OOD - 40% |  |
| | PCC ↑ | $R^2$ ↑ | PCC ↑ | $R^2$ ↑ | PCC ↑ | $R^2$ ↑ | PCC ↑ | $R^2$ ↑ | PCC ↑ | $R^2$ ↑ |
| DLKcat | 0.538±0.149 | 0.273±0.166 | 0.454±0.057 | 0.101±0.185 | 0.379±0.086 | 0.009±0.188 | 0.270±0.080 | -0.113±0.199 | 0.116±0.035 | -0.268±0.100 |
| UniKP | 0.773±0.002 | 0.578±0.004 | 0.590±0.026 | 0.271±0.020 | 0.567±0.036 | 0.246±0.023 | 0.503±0.042 | 0.184±0.024 | 0.424±0.062 | 0.112±0.029 |
| CatPred | 0.763±0.003 | 0.578±0.004 | 0.643±0.014 | 0.413±0.018 | 0.579±0.025 | 0.333±0.032 | 0.513±0.037 | 0.256±0.040 | 0.422±0.059 | 0.168±0.047 |
| OmniESI | 0.821±0.002 | 0.673±0.003 | 0.666±0.005 | 0.429±0.010 | 0.647±0.006 | 0.398±0.012 | 0.520±0.010 | 0.232±0.017 | 0.402±0.020 | 0.120±0.022 |
| CatESO | 0.826±0.002 | 0.680±0.004 | 0.671±0.010 | 0.436±0.018 | 0.663±0.011 | 0.416±0.019 | 0.553±0.019 | 0.257±0.027 | 0.514±0.024 | 0.189±0.031 |

| Medium |  |  |  |  |  |  |  |  |  |  |
| --- | --- | --- | --- | --- | --- | --- | --- | --- | --- | --- |
| Model | ID - 100% |  | OOD - 99% |  | OOD - 80% |  | OOD - 60% |  | OOD - 40% |  |
| | PCC ↑ | $R^2$ ↑ | PCC ↑ | $R^2$ ↑ | PCC ↑ | $R^2$ ↑ | PCC ↑ | $R^2$ ↑ | PCC ↑ | $R^2$ ↑ |
| DLKcat | 0.558±0.126 | 0.289±0.144 | 0.462±0.094 | 0.165±0.152 | 0.426±0.121 | 0.129±0.157 | 0.417±0.118 | 0.134±0.125 | 0.457±0.122 | 0.162±0.142 |
| UniKP | 0.749±0.002 | 0.556±0.002 | 0.590±0.009 | 0.337±0.009 | 0.571±0.008 | 0.299±0.008 | 0.579±0.016 | 0.287±0.011 | 0.542±0.020 | 0.257±0.014 |
| CatPred | 0.775±0.001 | 0.597±0.002 | 0.618±0.009 | 0.376±0.012 | 0.606±0.008 | 0.367±0.009 | 0.609±0.014 | 0.366±0.016 | 0.664±0.006 | 0.424±0.011 |
| OmniESI | 0.776±0.002 | 0.597±0.004 | 0.631±0.013 | 0.387±0.021 | 0.627±0.015 | 0.381±0.022 | 0.636±0.014 | 0.390±0.021 | 0.627±0.022 | 0.379±0.030 |
| CatESO | 0.782±0.001 | 0.607±0.002 | 0.640±0.002 | 0.401±0.003 | 0.640±0.004 | 0.398±0.008 | 0.652±0.006 | 0.410±0.008 | 0.670±0.005 | 0.424±0.006 |

| Long |  |  |  |  |  |  |  |  |  |  |
| --- | --- | --- | --- | --- | --- | --- | --- | --- | --- | --- |
| Model | ID - 100% |  | OOD - 99% |  | OOD - 80% |  | OOD - 60% |  | OOD - 40% |  |
| | PCC ↑ | $R^2$ ↑ | PCC ↑ | $R^2$ ↑ | PCC ↑ | $R^2$ ↑ | PCC ↑ | $R^2$ ↑ | PCC ↑ | $R^2$ ↑ |
| DLKcat | 0.511±0.178 | 0.259±0.207 | 0.480±0.009 | 0.171±0.024 | 0.476±0.010 | 0.173±0.018 | 0.495±0.035 | 0.184±0.060 | 0.575±0.040 | 0.248±0.056 |
| UniKP | 0.755±0.010 | 0.541±0.013 | 0.544±0.037 | 0.191±0.034 | 0.536±0.043 | 0.167±0.037 | 0.475±0.051 | 0.122±0.042 | 0.389±0.053 | 0.094±0.034 |
| CatPred | 0.757±0.004 | 0.563±0.006 | 0.589±0.018 | 0.342±0.024 | 0.610±0.023 | 0.367±0.026 | 0.566±0.028 | 0.317±0.032 | 0.576±0.030 | 0.308±0.041 |
| OmniESI | 0.789±0.005 | 0.619±0.007 | 0.644±0.017 | 0.397±0.018 | 0.643±0.019 | 0.391±0.019 | 0.546±0.028 | 0.265±0.026 | 0.474±0.037 | 0.201±0.033 |
| CatESO | 0.805±0.003 | 0.646±0.005 | 0.655±0.016 | 0.408±0.013 | 0.661±0.014 | 0.411±0.011 | 0.558±0.017 | 0.276±0.016 | 0.497±0.022 | 0.213±0.015 |

**Table S4:** Single-model comparison of CatESO fusion variants and baseline models for ID and OOD *k*_cat_ prediction. The upper panel reports PCC and MAE, and the lower panel reports RMSE and *R*^2^. Test splits were defined by the maximum sequence identity to the training set, with 100% used for ID evaluation and 99%, 80%, 60% and 40% for progressively stricter OOD evaluation. CF denotes concat fusion, and SF denotes simple fusion.

| Model | ID - 100% |  | OOD - 99% |  | OOD - 80% |  | OOD - 60% |  | OOD - 40% |  |
| --- | --- | --- | --- | --- | --- | --- | --- | --- | --- | --- |
| | PCC ↑ | MAE ↓ | PCC ↑ | MAE ↓ | PCC ↑ | MAE ↓ | PCC ↑ | MAE ↓ | PCC ↑ | MAE ↓ |
| CatPred | 0.748±0.005 | 0.754±0.007 | 0.552±0.024 | 1.107±0.031 | 0.523±0.028 | 1.155±0.036 | 0.483±0.039 | 1.259±0.043 | 0.473±0.053 | 1.266±0.059 |
| OmniESI | 0.765±0.006 | 0.758±0.009 | 0.606±0.016 | 1.036±0.019 | 0.594±0.019 | 1.065±0.021 | 0.556±0.024 | 1.155±0.025 | 0.495±0.033 | 1.211±0.030 |
| CF/CatESO | 0.783±0.004 | 0.709±0.007 | 0.629±0.014 | 1.000±0.020 | 0.623±0.016 | 1.026±0.022 | 0.590±0.022 | 1.121±0.026 | 0.555±0.031 | 1.163±0.034 |
| SF/CatESO | 0.781±0.005 | 0.711±0.012 | 0.625±0.014 | 1.001±0.020 | 0.619±0.016 | 1.030±0.024 | 0.585±0.020 | 1.126±0.025 | 0.551±0.026 | 1.163±0.035 |
| | RMSE ↓ | $R^2$ ↑ | RMSE ↓ | $R^2$ ↑ | RMSE ↓ | $R^2$ ↑ | RMSE ↓ | $R^2$ ↑ | RMSE ↓ | $R^2$ ↑ |
| CatPred | 1.093±0.010 | 0.555±0.008 | 1.483±0.036 | 0.265±0.035 | 1.529±0.041 | 0.234±0.041 | 1.628±0.052 | 0.197±0.050 | 1.596±0.067 | 0.175±0.069 |
| OmniESI | 1.062±0.012 | 0.575±0.009 | 1.395±0.026 | 0.347±0.024 | 1.413±0.029 | 0.335±0.028 | 1.506±0.032 | 0.292±0.030 | 1.554±0.038 | 0.222±0.038 |
| CF/CatESO | 1.023±0.009 | 0.605±0.007 | 1.357±0.023 | 0.382±0.021 | 1.367±0.025 | 0.378±0.023 | 1.457±0.031 | 0.336±0.028 | 1.482±0.038 | 0.292±0.036 |
| SF/CatESO | 1.027±0.011 | 0.603±0.009 | 1.363±0.023 | 0.376±0.021 | 1.374±0.027 | 0.371±0.025 | 1.466±0.030 | 0.329±0.027 | 1.489±0.034 | 0.286±0.033 |

**Figure S1:**
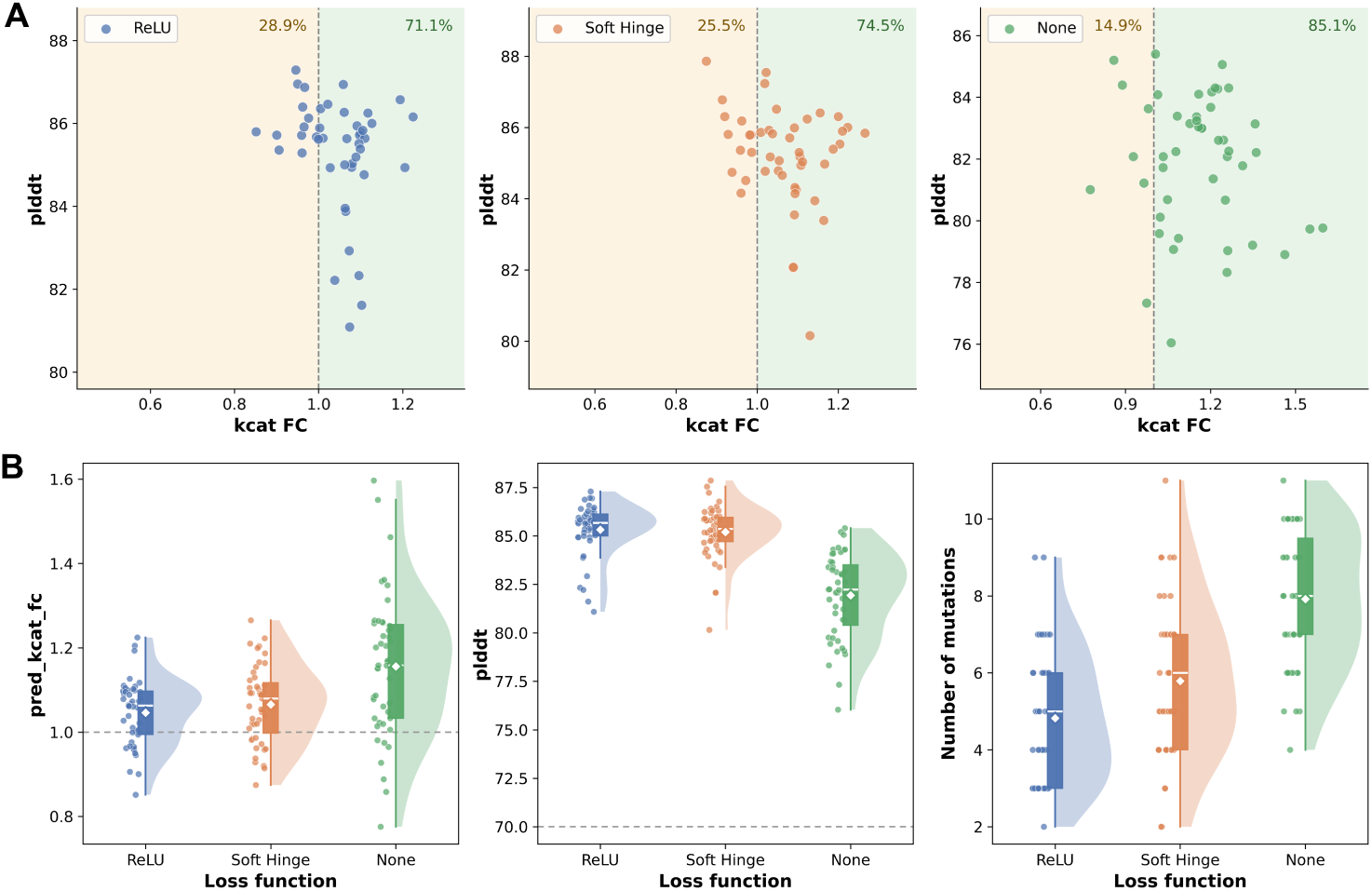
Cross-model validation of optimized Q9P9E3 sequences under different structural penalties. Fifty Q9P9E3 sequences were optimized under a ReLU, soft-hinge, or no (None) structural penalty, then re-scored with two evaluators held out from optimization: Protenix for structural confidence (pLDDT) and OmniESI for predicted activity (*k_cat_* fold change). (A) Per-sequence pLDDT against *k_cat_* fold change for the three penalty settings. Dashed lines mark *k_cat_* FC = 1 and the pLDDT threshold; percentages give the fraction of sequences on each side of the FC cutoff. (B) Distributions of OmniESI *k*_cat_ fold change, Protenix pLDDT, and number of mutations for the three settings. Box plots show the median and interquartile range; the grey dashed line marks the wild-type baseline (FC = 1) and the pLDDT reference at 70.

**Figure S2:**
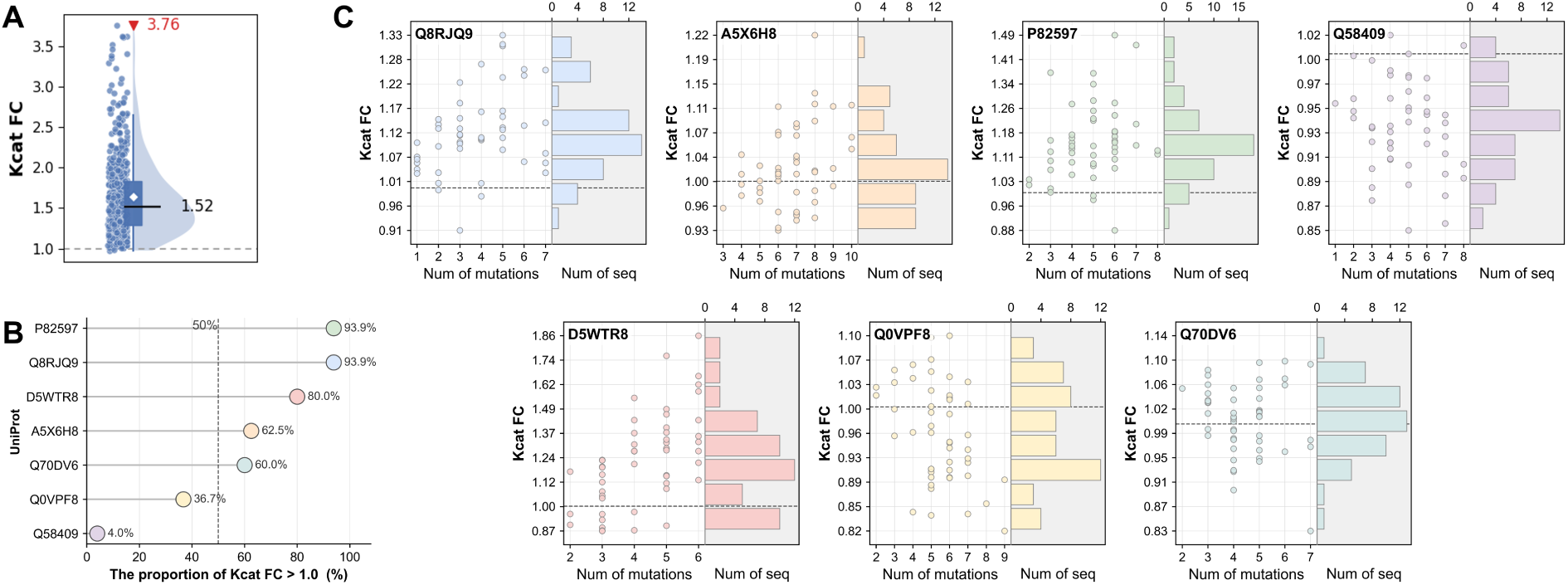
Predicted activity gains of CatESO-optimized variants across seven OOD enzymes. (A) CatESO-predicted *k*_cat_ fold-change distribution for non-redundant optimized variants, normalized to the corresponding wild-type sequence. (B) Fraction of beneficial variants, defined as optimized designs with predicted *k*_cat_ fold change *>* 1, for each OOD enzyme. (C) OmniESI-predicted *k*_cat_ fold changes for the optimized variants. The scatter plot shows fold change versus the number of substitutions, and the marginal histogram shows sequence counts.

**Figure S3:**
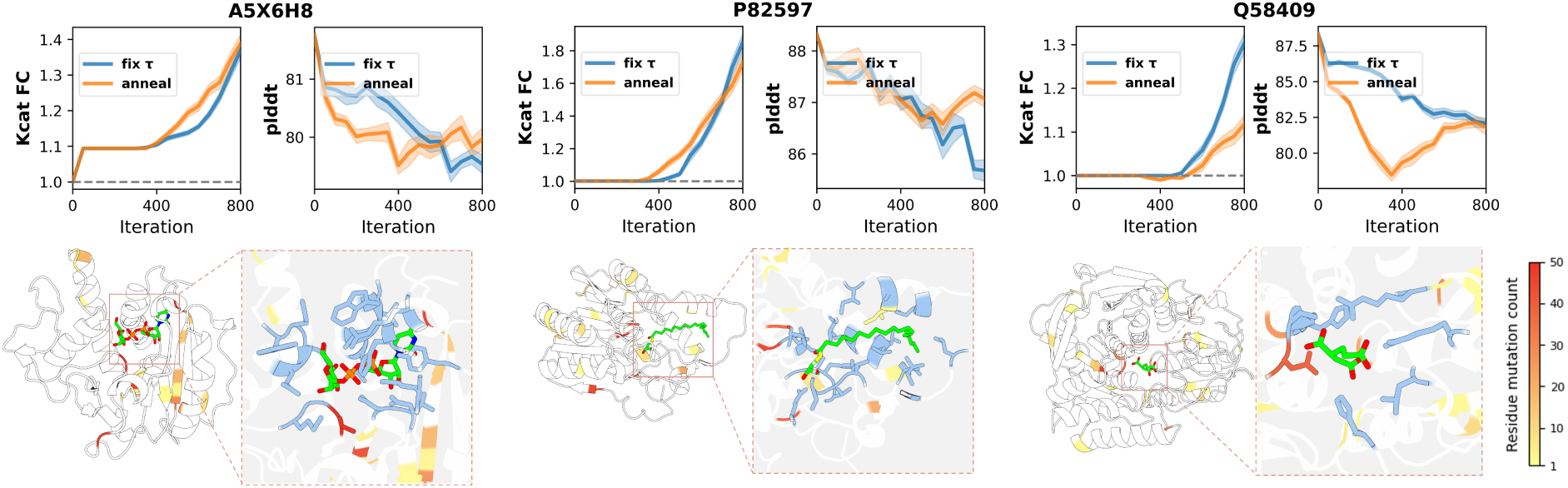
Additional per-enzyme optimization and structural analysis. Per-enzyme optimization and structural analysis for A5X6H8, P82597 and Q58409. Top panels show trajectories of CatESO-predicted *k*_cat_ fold change and of mean ESMFold pLDDT for the structurally relaxed sequences under the fixed (*τ* = *τ*_0_) and annealed (*τ* ! ↓) temperature schedules; solid lines denote the mean, shaded regions the ± s.d., and grey dashed lines the wild-type baseline. Bottom panels map CatESO-selected mutation frequencies onto a cartoon of the full enzyme structure and a magnified active-site view, with active-pocket residues shown in blue and color intensity scaled to per-residue mutation frequency.

**Figure S4:**
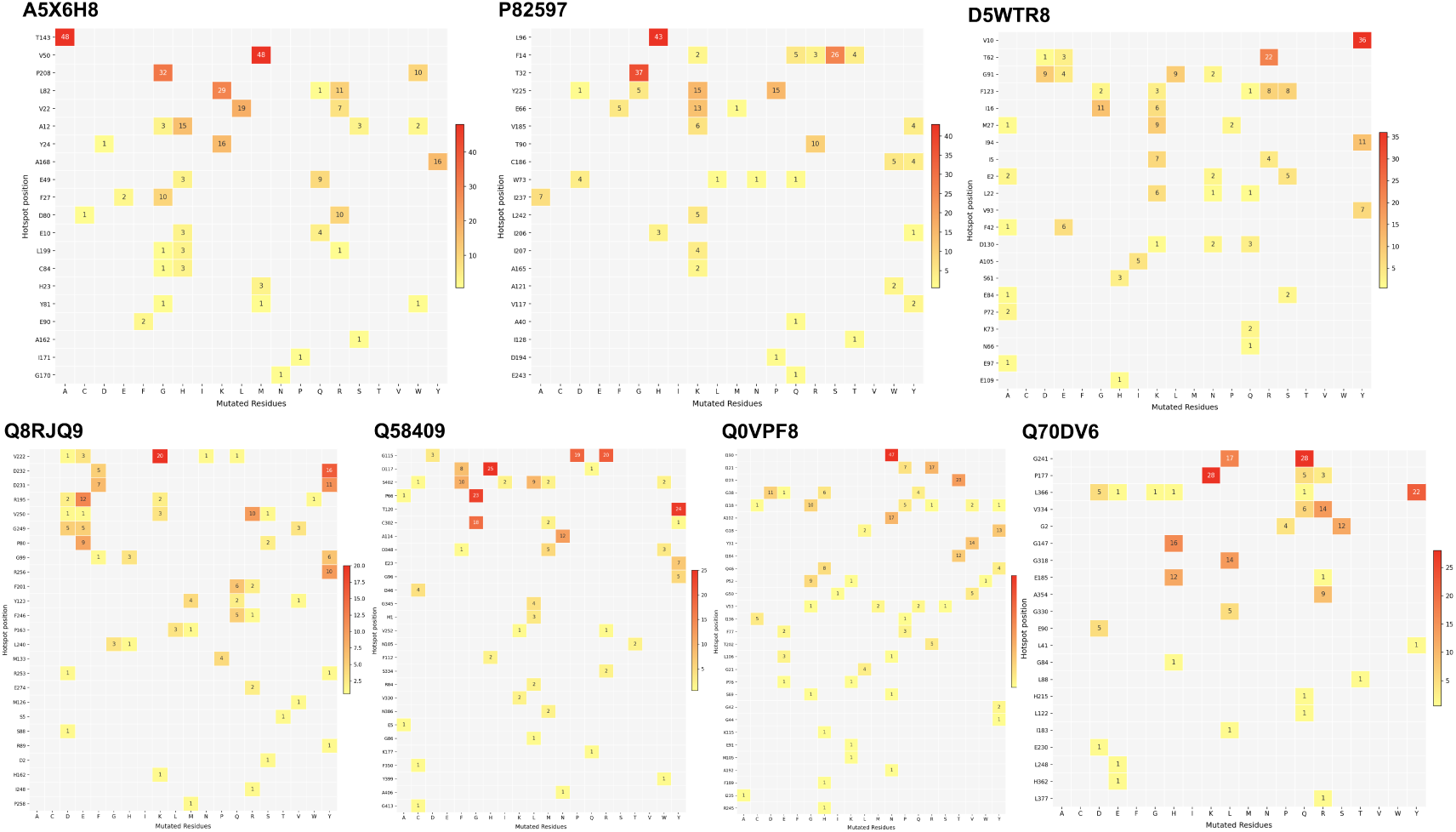
Mutational hotspot landscapes of CatESO-optimized variants across seven OOD enzymes. For each OOD enzyme, heatmaps summarize substitutions across non-redundant optimized variants. Rows correspond to wild-type residue positions and columns to substituted amino-acid identities. Colour intensity indicates substitution frequency, with rows ranked by total frequency to identify recurrently targeted positions.

**Figure S5:**
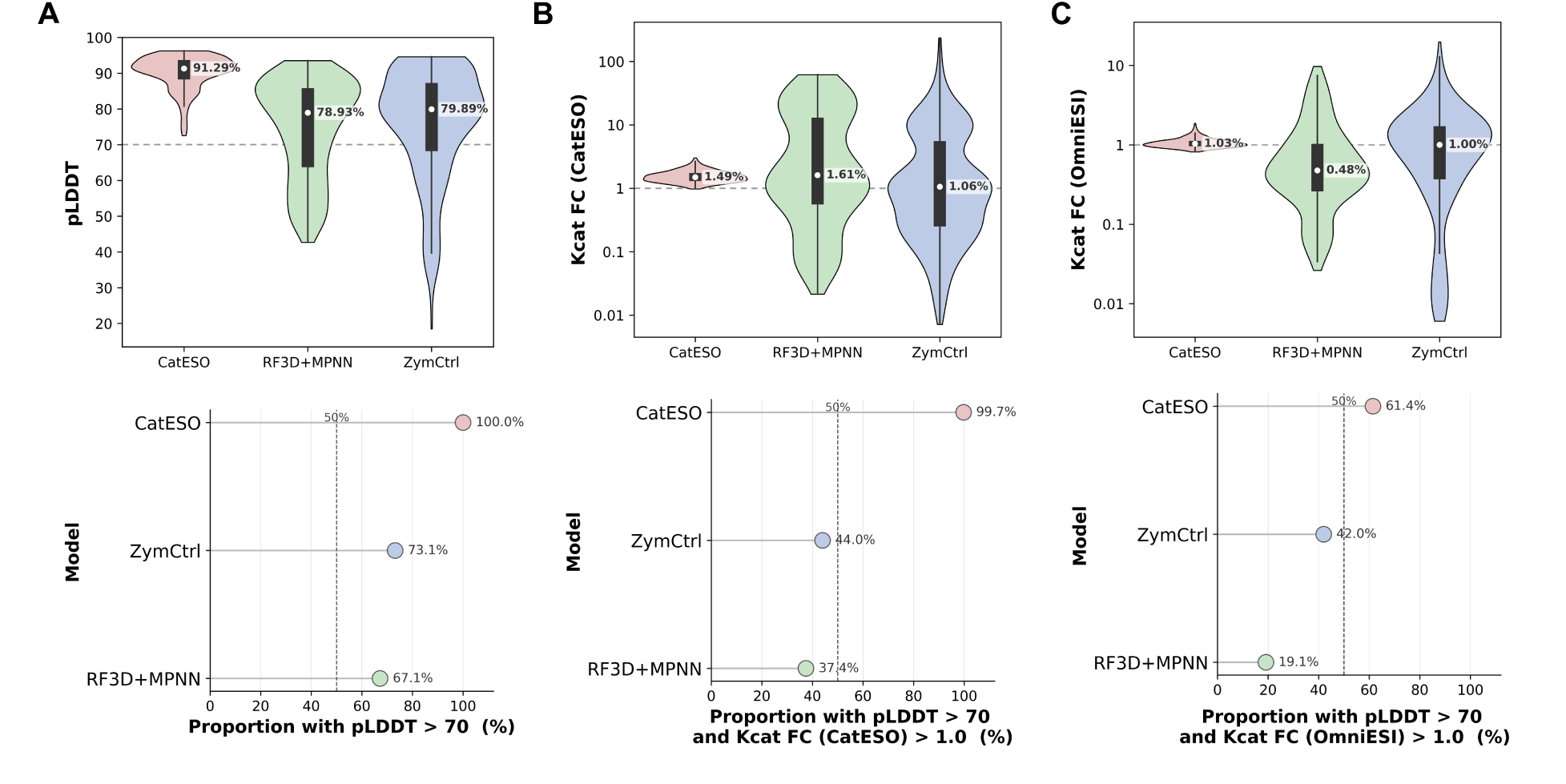
Predicted catalytic and structural quality of variants generated by CatESO, RF3D+MPNN and ZymCtrl. (A) Top: pLDDT distributions for each method; the dashed line marks the confidence threshold. In all violin plots, the black line denotes the median and the white diamond the mean. Bottom: fraction of structurally confident variants per method, defined by pLDDT ¿ 70. (B) Top: distributions of predicted *k_cat_* fold change for non-redundant variants, normalized to the corresponding wild-type enzyme and scored by CatESO. Bottom: fraction of variants meeting both the structural and activity criteria—high pLDDT and predicted *k_cat_* improvement over wild type—under CatESO scoring. (C) As in (B), but scored by OmniESI.

